# Synthetic Zippers as an Enabling Tool for Engineering of Non-Ribosomal Peptide Synthetases

**DOI:** 10.1101/2020.05.06.080655

**Authors:** Kenan A. J. Bozhueyuek, Jonas Watzel, Nadya Abbood, Helge B. Bode

## Abstract

Non-ribosomal peptide synthetases (NRPSs) are the origin of a wide range of natural products, including many clinically used drugs. Engineering of these often giant biosynthetic machineries to produce novel non-ribosomal peptides (NRPs) at high titre is an ongoing challenge. Here we describe a strategy to functionally combine NRPS fragments of Gram-negative and -positive origin, synthesising novel peptides at titres up to 290 mg l^-1^. Extending from the recently introduced definition of e*X*change *U*nits (XUs), we inserted synthetic zippers (SZs) to split single protein NRPSs into up to three independently expressed and translated polypeptide chains. These synthetic type of NRPS (type S) enables easier access to engineering, overcomes cloning limitations, and provides a simple and rapid approach to building peptide libraries via the combination of different NRPS subunits.

**One Sentence Summary:** Divide and Conquer: A molecular tool kit to reprogram the biosynthesis of non-ribosomal peptides.

## Introduction

Non-ribosomal peptide synthetases (NRPSs) are multifunctional enzymes, producing a broad range of structural diverse and valuable compounds with diverse applications in medicine and agriculture (*1*) making them key targets for bioengineering. The structural diversity of non-ribosomal peptides (NRPs) arises from the assembly line architecture of their biosynthesis. According to their biosynthetic logic, known NRPS systems are classified into three groups, linear (type A), iterative (type B), and nonlinear NRPSs (type C) (*2*). Type A NRPSs are composed of sequential catalytically active domains organised in modules, each responsible for the incorporation and modification of one specific amino acid (AA). The catalytic activity of a canonical module is based upon the orchestrated interplay of an adenylation (A) domain for AA selection and activation, a condensation (C) domain to catalyse peptide bond formation, and a thiolation/ peptidyl-carrier protein (T) onto which the AAs or intermediates are covalently tethered (*3*). In addition, tailoring domains, including epimerization (E), methylation, and oxidation domains can be part of a module, or a heterocyclization (Cy) domain instead of a C-domain can be present. Finally, most NRPS termination modules harbour a TE-domain, usually responsible for the release of linear, cyclic or branched cyclic peptides (*4*).

Type A NRPSs (Fig. 1a) follow the collinearity rule, *i.e.* the number of NRPS modules corresponds directly to the number of monomers incorporated into the associated product, and the arrangement of the modules directly follows the peptides primary sequence (*5*). Whereas in *in cis* type A NRPSs all modules are arranged on a single polypeptide chain (*e.g.* ACV-synthetase (*6*)), *in trans* assembly-lines comprise a number of individual proteins (Daptomycin-synthetase (*7*)). Mutual protein-protein interactions of the latter are mediated by specialized *C*-(donor) and *N*-terminally (acceptor) attached ∼30 AAs long *α*-helical structural elements, so called communication mediating (COM) or docking domains (DDs) (*8*). DDs typically are located in between two modules and only interact with weak affinities (4-25 µM) (*9–13*), but are crucial to ensure biosynthesis of the desired product(s) (*8, 11, 14*). Despite recent progress on applying DD substitutions to program new assembly lines, in most cases structural information is lacking to effectively apply DDs for general engineering purposes (*11, 15, 16*).

**Figure 1.**
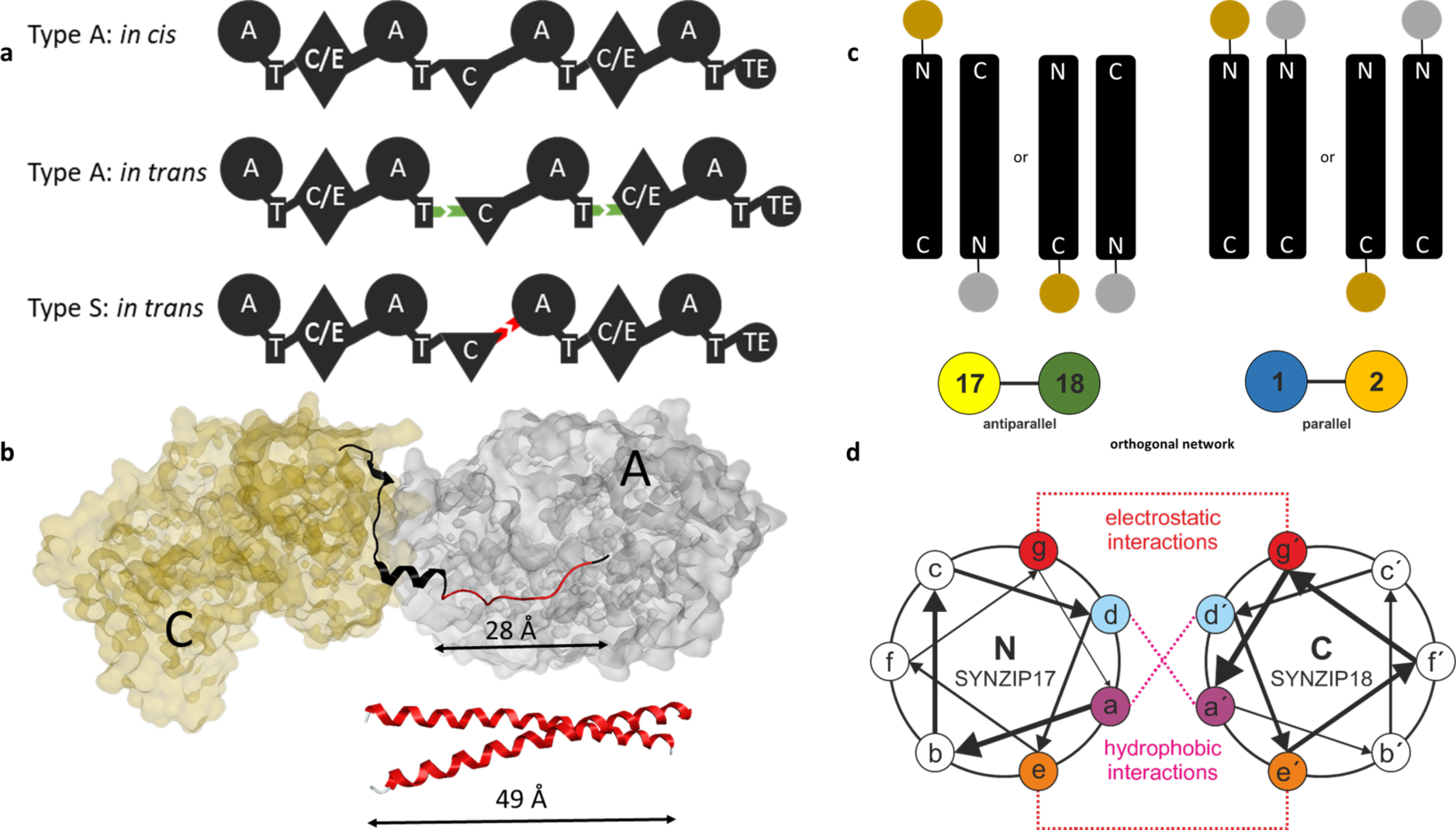
Introduction to SZ mediated in trans protein-protein interaction. **(a)** Schematic overview of type A and type S NRPSs. Natural DDs (*in trans* type A NRPS) are shown in green and artificial SZs in red (*in trans* type S NRPS). For domain assignment the following symbols are used: A, adenylation domain, large circles; T, thiolation domain, rectangle; C, condensation domain, triangle; C/E, dual condensation/epimerization domain, diamond; TE, thioesterase domain, small circle. **(b)** Top: excised C-A di-domain and linker region (ribbon representation) from the SrfA-C termination module (PDB-ID: 2VSQ). Removed area of the C-A linker region to introduce SZs is highlighted red. Bottom: modelled 41 AAs comprising SZ pair. **(c)** Top: antiparallel (left) and parallel (right) interacting hetero-specific SZs. Bottom: SZ17:18 and SZ1:2 are forming an orthogonal interaction network. **(d)** SZ interactions. SZ17:18 are predicted to be electrostatic complementary at adjacent interfacial *e* and *g* positions.

Although early engineering attempts, including the exchange of DDs, the targeted modification of the A-domains substrate specificity conferring AA residues, and the substitution of domains as well as whole modules, gave mixed results, several notable advances have been published recently (*16–18*). To give but one example, we comprehensively analysed structural data as well as inter-domain linkers in NRPSs to define novel fusion sites and to provide guidelines for exchanging A-T-C units, denoted as e*X*change *U*nits (XUs), as opposed to canonical modules (C-A-T) (*19*). By combining XUs from 15 NRPSs *in cis*, it was possible to reconstitute naturally available peptides, peptide derivatives, and to generate new-to-nature peptides *de novo* in high yields.

Herein, starting from our recently published XU concept, we explored the ability of synthetic zippers (SZs (*20*)) to manipulate collinear type A NRPSs by introducing artificial *in trans* regulation. SZs interact with high affinity (KD<10 nM) via a coiled-coil structural motif, enabling the specific association of two proteins. Such a strategy not only would allow creating a synthetic type of *in trans* regulated mega-synthetases (type S), by combining NRPSs with high-affinity SZs (*20*) (Fig. 1a), but to overcome cloning and protein size limitations associated with heterologous NRP production.

## Results

In depth structural analysis of the crystallised termination module SrfA-C (PDB-ID: 2VSQ) suggested splitting NRPSs in between consecutive XUs at the previously defined W]-[NATE motif of the conformationally flexible C-A linker (*21–23*) region. As already known (*21, 22*), this splicing position bears several advantages. Of particular importance is that it keeps intact the short (∼ 10 AAs) *α*-helical structure at the *C*-terminus of the resulting truncated protein (**subunit 1**) – as in wild type (WT) NRPSs this helical structure not only regulates the C-A distance throughout the catalytic cycle (*21*), but also associates with the A-domains hydrophobic protein surface (*23*).

Attempts of *in silico* creating NRPS domains connected via SZs, composed of ∼40 amino acids (AAs), were unsuccessful. Nevertheless, careful revision of available structural data indicated that ∼10 AAs from the unstructured *N*-terminus of **subunits 2** must be removed to meet the distance-criteria set out by the WT C-A inter-domain linker to ensure correct C-A di-domain contacts before SZs *N*-terminally could be introduced (Fig. 1b). After perusing characterized SZ pairs (*20*), to begin with we chose the SZ pair 17 & 18 (Fig. 1c & d).

### Proof of Concept

To assess the general suitability of SZ-pairs to *in trans* connect two NRPS proteins and mediate biosynthetically functional protein-protein interface interactions, we targeted the xenotetrapeptide (**1**) producing NRPS (XtpS; Fig. S1) from the Gram-negative entomopathogenic bacterium *Xenorhabdus nematophila* HGB081 (*24*). We decided to split XtpS into two subunits in between XUs 2 and 3 and four artificial two component type S NRPS (Fig. 2a) were constructed and heterologously produced in *E. coli* DH10B::*mtaA* (*25*) – either with SZs fused to both subunits (NRPS-1: subunit 1-SZ17, SZ18-subunit 2); only fused to subunit 1 (NRPS-2: subunit 1-SZ17, subunit 2) or subunit 2 (NRPS-3: subunit 1, SZ18-subunit 2), and without SZs (NRPS-4: subunit 1, subunit 2).

**Figure 2.**
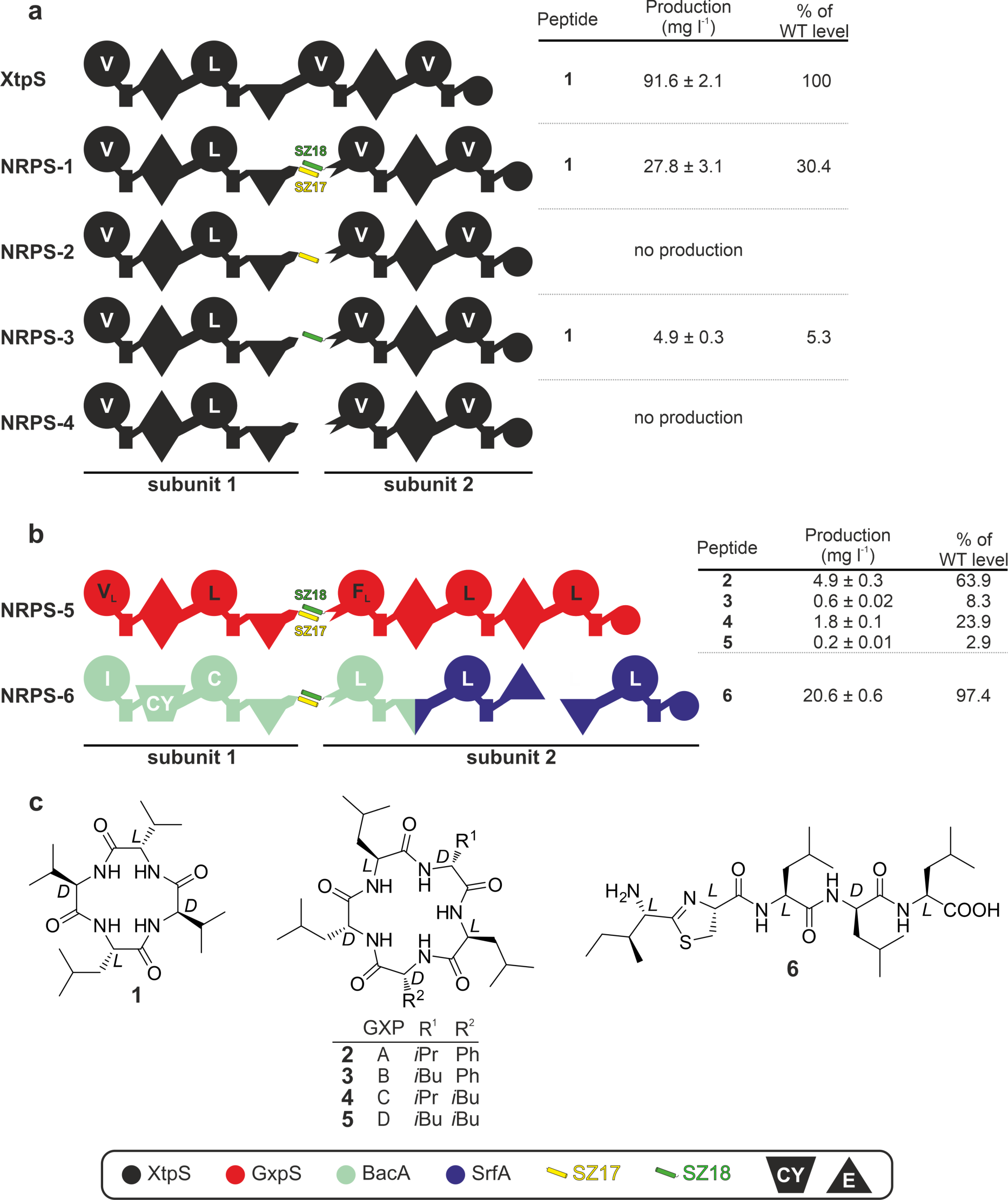
**(a)** Type S **NRPS-1** – **4**, as well as corresponding peptide yields obtained from triplicate experiments. **(b)** Type S **NRPS-5** and **-6**, where GxpS and RtpS are split in two subunits. **(c)** Structures of **1**–**6** produced from **NRPS-1** to **NRPS-6** expressed in *E. coli*. See Fig. 1 for assignment of the domain symbols; further symbols: CY, heterocyclization domain; E, epimerization domain. Boxed are the colour coded NRPSs used as building blocks and the used SZ pairs.

NRPS-2 and NRPS-4 showed no detectable peptide production, whereas NRPS-1 lead to the production of **1** with ∼30% (28 mg l^-1^) yield compared to WT XtpS (Fig. 2a, Fig. S2), confirming that SZs indeed can be used to functionally mediate new-to-nature *in trans* regulation of NRP biosynthesis. Interestingly, NRPS-3 with SZ18 fused to subunit 2, but lacking SZ17 on subunit 1, showed moderate yields of **1**. Despite lacking SZ17, the *C*-terminus of XtpS subunit 1 is forming a Leucine rich *α*-helical structure (PDB-ID: 2VSQ) that might be able to interact with SZ18 of subunit 2 and mediate an impaired but catalytically active C-A interface (*21–23, 26*).

Additionally, SZ17:18 were used to split the GameXPeptide A-D (**2**-**5**) producing NRPS (GxpS (*27, 28*)) and the recombinant thiazole-peptide (**6**) producing NRPS (RtpS (*29*)). Whereas GxpS originates from the Gram-negative bacterium *Photorhabdus luminescens* TTO1, RtpS was constructed previously (*29*) from building blocks (BBs) of Gram-positive origin (using NRPSs for the production of bacitracin (*30*) and surfactin (*31*)). Both resulting type S NRPSs (Fig. 2b) showed good to very good titres of desired peptides. NRPS-5 produced **2** (Fig. S3) with yields of ∼64 % (4.9 mg l^-1^) compared to WT GxpS and NRPS-6 produced **6** (Fig. S4) at WT RtpS level (∼20 mg l^-1^).

All product structures and yields were confirmed by tandem mass spectrometry (MS/MS) analysis and comparison of the retention times with synthetic standards.

### Creating Synonymous Chimeras

To explore the recombination potential of chimeric type S NRPSs, initially we co-expressed non-cognate subunits from NRPS-1 and -5 (Fig. 3a). Both, NRPS-1 subunit 1 and NRPS-5 subunit 1, largely possess synonymous A-domain (Val, Leu) and C-domain (hydrophobic AAs) specificities, preventing potential upstream C- and downstream TE-domain substrate specificity issues (*22*).

**Figure 3.**
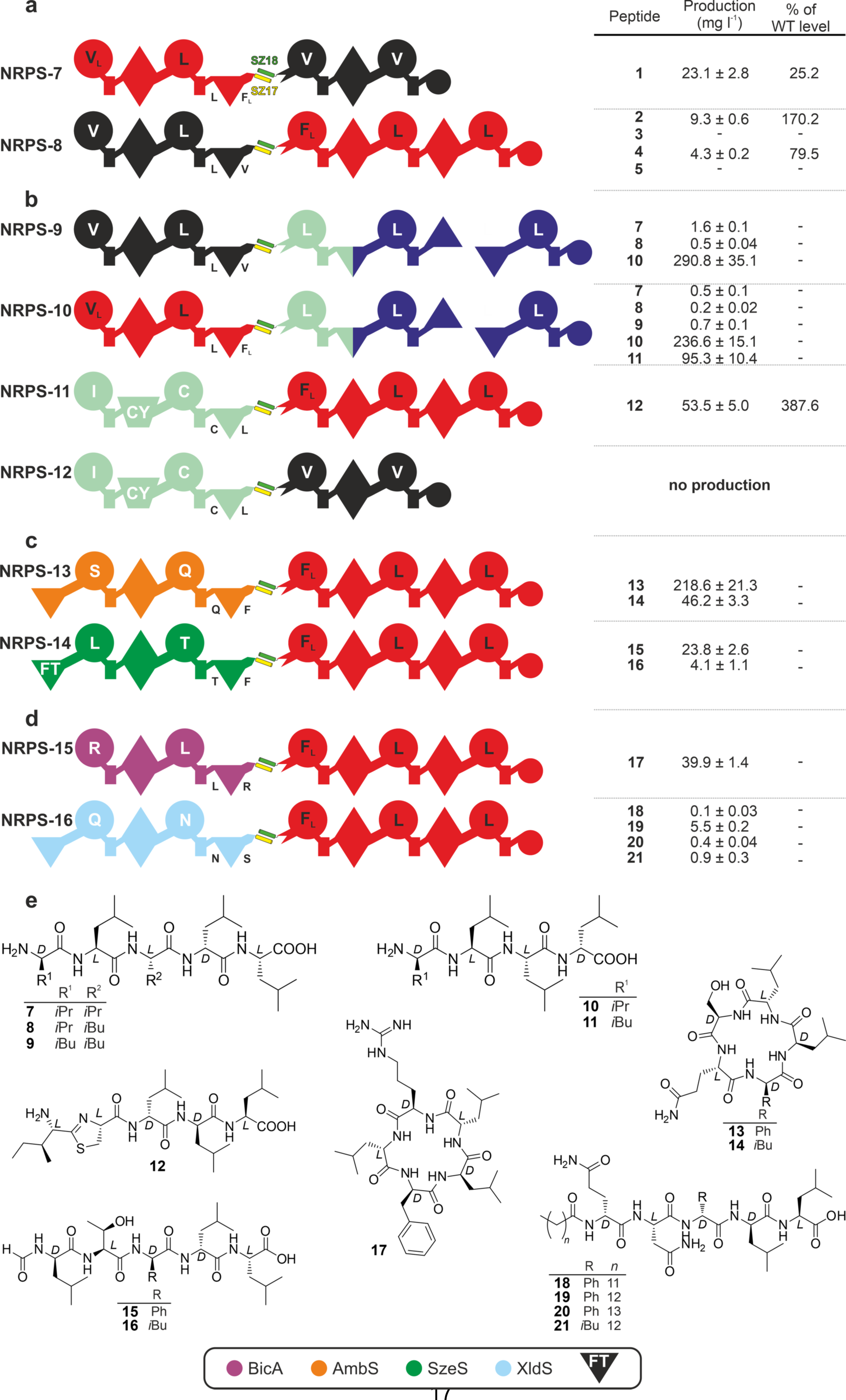
**(a)** Type S NRPSs using building blocks from Gram-negative bacteria (**NRPS-7** and **NRPS-8), (b)** from Gram-negative and -positive bacteria (**NRPS-9** – **NRPS-12), (c)** that consider the specificity of the C domain acceptor site (**NRPS-13** and **NRPS-14)** and **(d)** that do not consider the specificity of the C domain acceptor site (**NRPS-15** and **NRPS-16). (f)** The structures of **7**–**21** produced from **NRPS-9** to **NRPS-16** expressed in *E. coli*. See Fig. 1 and 2 for assignment of the domain symbols; further symbols: FT, formyl-transferase domain.

NRPS-7 produced **1** at same titres as NRPS-1 with 25.2 % (∼23 mg l^-1^) yield compared to WT XtpS (Fig. S5). NRPS-8 showed 4- to 5-fold increased productivity compared to NRPS-5 and even exceeded WT GxpS in yields of **2** and **4** with 170 % (∼9 mg l^-1^) and 80 %, respectively (Fig. S6). Like observed in our previous work (*22, 29*) the formal exchange of the promiscuous XU1 from GxpS (for Val/Leu) against the Val-specific XU1 from XtpS led to the exclusive production of **2** and **4**, without synthesis of **3** and **5** observed in the original GxpS. Increased peptide yields of NRPS-8 compared to its WT GxpS counterpart currently cannot be explained but were described before (*29*).

### Reprogramming Type S NRPS

Major drawbacks of past reprogramming attempts have been (I) the incompatibility of bacterial NRPS BBs from Gram-positive and -negative origin (*22, 29*) as well as (II) the C-domains’ specificity rule (*22*); both (I + II) severely limiting combinatorial space. With tools to functionally split NRPSs at hand, we explored if type S NRPS have the potential to overcome these biosynthetic bottlenecks.

To address (I), subunits of Gram-negative (NRPS-1 & -5) and -positive (NRPS-6) origin were co-expressed (Fig. 3b) and culture extracts analysed via HPLC-MS (Fig. S7-9). Three out of four resulting NRPSs (NRPS-9 – 12) not only showed detectable, but very good peptide titres up to ∼290 mg l^-1^ (NRPS-9, **10**). Yet, all active type S NRPSs showed unexpected peptide production profiles. For instance, NRPS-10 only produced trace levels of the expected linear penta-peptide (**8**, linear *v*LL*l*L; D-AAs in italics and lowercase throughout this work) but produced yields of ∼236 mg l^-1^ and ∼95 mg l^-1^ of linear *v*LL*I* (**10**) and *I*LL*I* (**11**), respectively. NRPS-9 and -10 not only constitute two polypeptide chains linked via SZs, but the termination-module also interacts *in trans* via natural DDs with subunit 2. Due to observed very high catalytic activities of NRPS-9 and -10, along with SZs interacting with much higher affinities than DDs, ‘auto-catalytic’ offloading might be catalysed by the E-domain present at the *C*-termini of respective subunits 2 as it was recently reported by non-cognate termination domains of chimeric NRPSs utilising internal C-domains (*22*).

When NRPS-11 and -12 were compared, the latter did not produce any detectable peptide amounts, likely being an issue of TE-domain substrate specificity of XtpS (*c.f.* Fig. S10), NRPS-11 produced the linear thiazole containing peptide IC\**ll*L (**12**; ∼53 mg l^-1^; Fig. S9). In its natural NRPS context as well as *in vitro*, the A3-domain of GxpS prefers Phe over Leu (*29*). In case of NRPS-11, the terminal C-domain of subunit 1, expecting Leu at its acceptor site, either prevents the incorporation of Phe due to its gatekeeping activity or rather fine tunes the downstream A-domain specificity. Similar effects of engineered NRPSs, exhibiting chimeric C-A interfaces (*22, 32*) or C-domains (*29*), have been described. Aforementioned results (NRPS-9 – 12, Fig. 3b) represent the first successful *in trans* as well as the first efficient strategy to recombine Gram-positive and -negative NRPS BBs, producing peptides at an industrially relevant scale.

Tackling (II), subunits of NRPSs with non-synonymous specificities were co-expressed with subunit 2 of NRPS-1 as well as NRPS-5, either respecting the C-domains’ specificity rule (NRPS-13 and 14, Fig. 3c) or not (NRPS-15 and 16, Fig. 3d; NRPS-17 – 19, Fig. S10). As subunits 1, XUs1-2 of NRPSs producing ambactin (AmbS (*25*)), szentiamide (SzeS (*22*)), bicornutin (BicA (*33*)), and xenolindicin (XldS (*25*)) were used.

Both type S NRPS (NRPS-13: AmbS subunit 1 + GxpS subunit 2; and NRPS-14: SzeS subunit 1 + GxpS subunit 2) respecting C-domains’ acceptor site specificities (Phe) produced the desired derivatives (Fig. 3c). NRPS-13 produced **13** and **14** with yields of ∼218 and ∼46 mg l^-1^, respectively (Fig. S11). In addition, peptide yields of **15** (∼24 mg l^-1^) and **16** (∼4 mg l^-1^) from NRPS-14 were even higher compared to a homologous *in cis* NRPS (**15**: ∼15 mg l^-1^; **16**: ∼2 mg l^-1^) constructed from the same BBs in an earlier study (*22*) (Fig. S12 & S13). Peptides **13**/**14** and **15**/**16**, respectively, only differ in Leu or Phe at position 3 from the relaxed substrate specificity of XU3 from GxpS (*27, 28*).

NRPS-15 (BicA subunit 1 + GxpS subunit 2) and -16 (XldS subunit 1 + GxpS subunit 2), not complying with the XUs’ specificity rules (Fig. 3d), produced peptides **17** (∼40 mg l^-1^, Fig. S14) and **18**-**21** (0.1 – 5.5 mg l^-1^, Fig. S15), respectively. The latter peptides (**18**-**21**) only differ in the *N*-terminal acyl starter unit, originating from the *E. coli* fatty-acid pool, as also observed in the original xenolindicins (*25*). Especially NRPS-15 was expected to be inactive, as previous studies have shown that the BicA C3-domains acceptor site is highly specific for Arg and cannot process Phe or Leu when covalently fused to subunit 2 (*29*). This might indicate that splitting *in cis* NRPSs in between C- and A-domains potentially decreases C-domains’ acceptor site specificity by introducing more geometric flexibility and minimizing potentially restrictive effects on A-domain movements (*32*) as supported by a recently published study suggesting that C-domains indeed do not exhibit intrinsic substrate specificities (*34*).

In contrast, all type S NRPS (NRPS-17 – 19, and NRPS-12) sharing subunit 2 from XtpS (NRPS-1), also not complying with the C-domain specificity rule (*22, 29*), failed to produce detectable amounts of any peptide (Fig. S10). The reason for this might be the TE-domains high specificity for peptide length and amino acid composition.

### Unpaired Activity of GxpS Subunit 2

All type S NRPS split in between C-A domains and sharing GxpS subunit 2 (NRPS-5, -8, -11, and NRPS-13 – 16) showed an unexpected behaviour, producing a range of tripeptides (**33/34** and **35/36**) at high titre up to 86 mg l^-1^ related to the unpaired activity of GxpS subunit 2 (Fig. S16). Due to the promiscuous GxpS A3-domain, **33/34** and **35/36** differ from each other at position one, either carrying Phenylalanine or Leucine. In addition, **33** and **35** show a *D*-*D*-*L* configuration, whereas **34** and **36** have a *L*-*D*-*L* configuration.

### Optimization and Extending Functionality

To explore the optimization potential of SZs we not only successfully applied parallel interacting (Fig. 1c) SZs 19 &18 (Fig. 4a & S17a, NRPS-20), but also optimized SZ17:18 interactions. Thus, to introduce more spatial freedom potentially enhancing flexible domain-domain interactions, synthetic stretches of Gly-Ser (GS) varying in length of 4-10 AAs were introduced in between the *C*-terminus of XtpS (NRPS-1) subunit 1 and SZ17 (Fig. 4b). All resulting chimeric NRPSs (NRPS-21 - 23) showed ∼3x increased yields of **1** (Fig. S17b) compared to NRPS-1, raising titres back at WT level (∼220 mg l^-1^) – and therefore indicating that introduced GS linkers had the desired effect.

**Figure 4.**
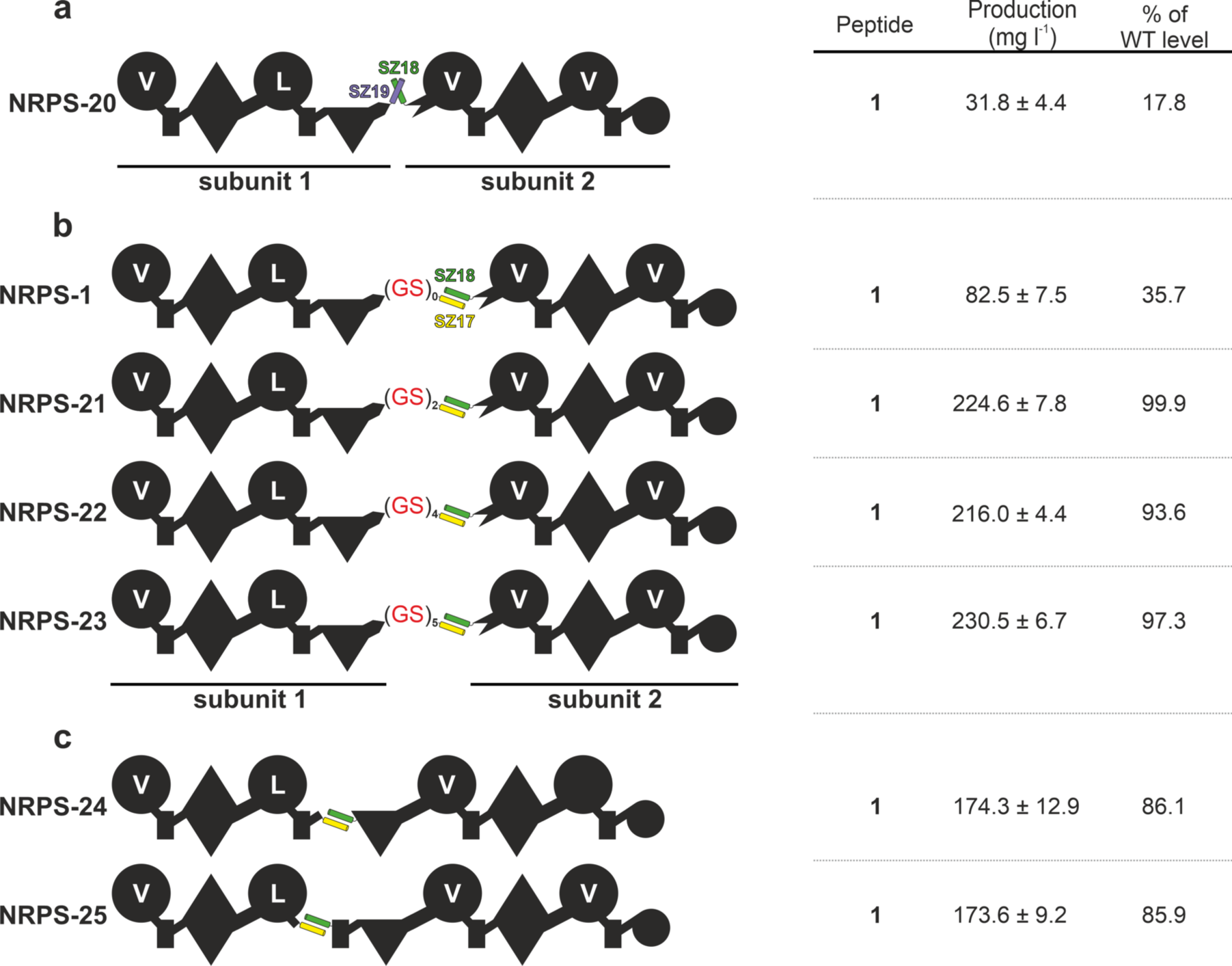
**(a)** Type S XtpS using parallel interacting SZ18:19, **(b)** (GS)_x_-elongated C-A linker sequences (**NRPS-21** – **NRPS-23), (c)** using different split positions in between the T-C (**NRPS-24**) and A-T domains (**NRPS-25**). See Fig. 1 for assignment of the domain symbols.

To further explore and extend functionality of type S NRPSs, we again targeted XtpS introducing SZs within the T-C (NRPS-24, Fig. S18a) and A-T linker-regions (NRPS-25, Fig. S18b). Both, NRPS-24 and -25 (Fig. 4c & S17c) synthesised **1** with titres at ∼86 % (174 mg l^-1^) compared to WT XtpS level. While catalytic activity of NRPS-24 was expected, as the introduced SZs are mimicking natural DDs (*35*), NRPS-25, showing an unusual split between A2-T2, truly represents a type S NRPS. The A-T linker sequence, consisting of ∼15 AAs, represents the shortest inter-domain linker in the context of NRPS elongation modules, which is conformationally also the most flexible one. Structural insights indicate that T and A_sub_ domains adopt alternative conformations to shuffle reaction intermediates among catalytic domains (*23*). Thus, it was expected that the additional rigidity, inserted by the structured *α*-helical AA stretches, would result in loss of function. Structures of large constructs of the linear gramicidin synthesising NRPS (LgrA) (*35*) show a very high structural flexibility, potentially bringing closely together domains that are far apart in protein sequence and therefore facilitating synthetic cycles with inserted tailoring domains, unusual domain arrangements like A-C-T (*36*), module skipping (*37*) and presumably also SZs.

More dipartite type S NRPSs (NRPS-44 - 48), split in between and within modules are depicted in Fig. S19 and S20.

### Tripartite NRPSs

SZs mediated reprogramming of NRPSs, other than ‘simply’ splitting *in cis* NRPSs in two, makes it necessary to express the proteins in three parts – *i.e.* being able to target one specific position of the synthesised peptide. Therefore, again starting with XtpS, we created three orthogonal interaction networks (Fig. 1c) by introducing the anti-parallel and parallel interacting SZ pairs SZ17:18 and SZ1:2 in between XU2-3 and XU3-4 (NRPS-26), in between module 2-3 and module 3-4 (NRPS-27), as well as within the A-T linker regions of module 2 and 3 (NRPS-28), respectively (Fig. 5).

**Figure 5.**
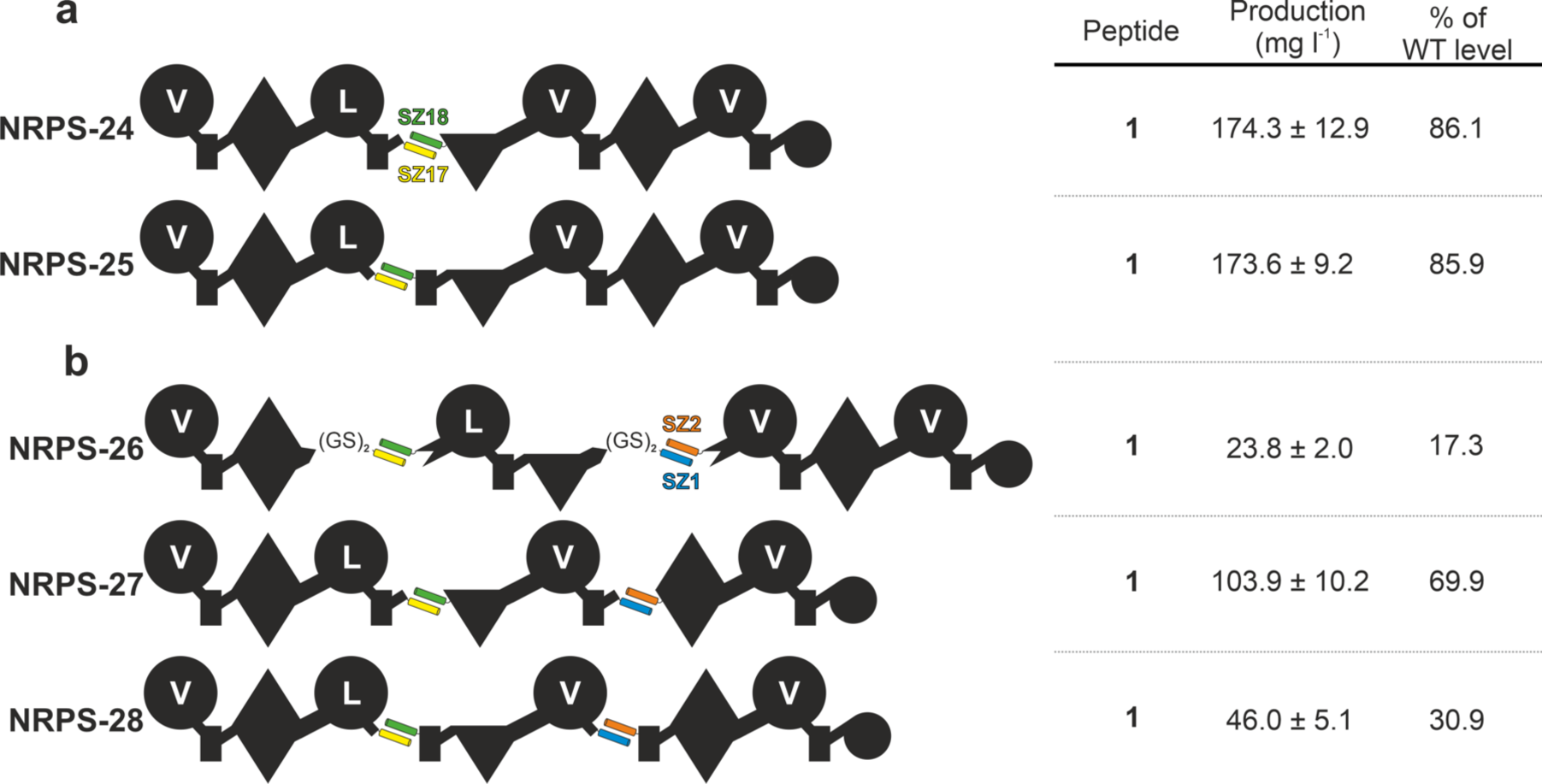
Schematic representation of three subunit type S XtpS using the SZ1:2 and SZ17:18 pairs splitting in between the C-A (**NRPS-26**), T-C (**NRPS-27**) and A-T linker region (**NRPS-28**). See Fig. 1 for assignment of the domain symbols.

All resulting tripartite type S NRPSs (NRPS-26 – 28, Fig. 5 & S17d) produced **1** with good yields of 17 – 70% (24 – 104 mgl^-1^) compared to WT XtpS but with decreased yields compared to their dipartite counterparts (NRPS-21, -24, & -25).

In order to demonstrate the potential of artificial *in trans* NRPSs, we designed and cloned a small library of type S BBs (Fig. 6a), placing SZ17:18 and SZ1:2 within the A-T linker regions to perform co-expression experiments (Fig. 6) in a quick plug-and-play manner. The A-T linker region was targeted because in this case C-domain specificities presumably do not represent a limitation of recombination and because T-C-A tri-domains as a catalytically active unit to reprogram NRPSs are under underrepresented (*38, 39*).

**Figure 6.**
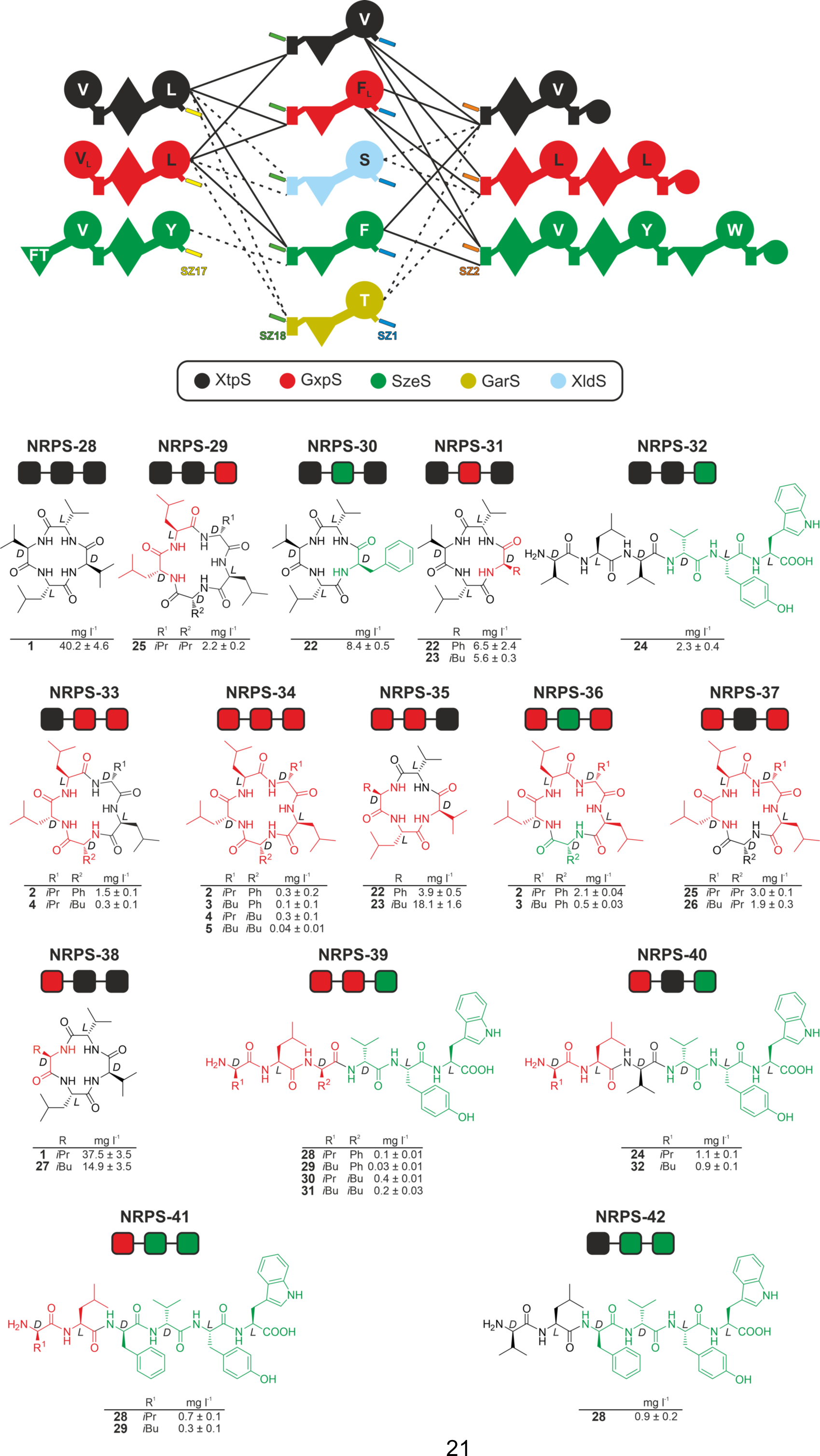
Schematic representation of recombinant type S NRPSs based on three subunit combinations (**NRPS-28** – **NRPS-42**) using the A-T linker region as split position leading to a structurally diverse peptide library of **22**–**32** all expressed in *E. coli*. See Fig. 1 and 3 for assignment of the domain symbols.

In brief, the created plasmid library, expressing 11 different type S NRPS BBs from XtpS, GxpS, SzeS, XldS, and the gargantuanin producing synthetase (GarS). Overall 15 (NRPS-28 – 42) from 22 co-expressions of three plasmids each yielded detectable amounts (0.1 – 38 mg l^-1^) of 16 different peptides, 11 of which were new (Fig. 6, S21 – S38). Despite the method’s general simplicity, the overall efficacy or recombination potential of T-C-A units compared to XUs appears to be slightly more restricted. For example, neither co-expression of all type S BBs to reconstitute SzeS, nor any combination involving the Ser and Thr specifying BBs from XldS and GarS, yielded any detectable peptide, respectively. These results probably indicate an incompatibility of formed chimeric A-T interfaces or substrate incompatibilities at the respective C-domains donor site. Yet, in light of previous results (*40, 41*) concerning C-domain specificities, the latter seems to be unlikely.

## Conclusion

Recently the successful application of SZs to replace naturally present DDs in polyketide synthases (PKS) as a tool to create chimeric PKSs was published (*42*). Here we reported the use of high-affinity SZs to split native single protein NRPSs into two and three individual components. Generating artificially *in trans* regulated assembly lines not only represents a new NRPS architecture, referred to as type S, but emerges to be highly productive with yields comparable to WT levels. Especially the efficient combination of NRPS BBs of Gram-negative and -positive origin greatly expands the combinatorial space for reprogramming pharmaceutically relevant entities or creating diverse NRP libraries.

Observed accumulation of side products may be a drawback, especially for industrial production purposes, but also indicates the high productivity potential of type S NRPS. For example, aggregated peptide yields of tetra- and penta-peptides produced by NRPS-10 sums up to ∼330 mg l^-1^, whereas the WT counterparts in total “only” produced ∼12 mg l^-1^ (GxpS) and ∼22 mg l^-1^ (RtpS), respectively. Increased total peptide yields, *i.e.* for type S NRPS split in between XUs, might be due to higher catalytic activities of C- and A-domains involved in forming the SZs mediated chimeric C-A interface, as geometric restrictions and feedback mechanisms of the catalytic cycle may have been suppressed.

Although it was not possible to insert SZs in between the *N*- and *C*-lobe of C-domains (Fig. S19c), following our recently published XUC concept (*29*), in principle it was possible to introduce SZs in between any di-domain (A-T, C-A, T-C). Having SZs at hand, not only peptide libraries quickly can be constructed with high success rates, but now it also should be conceivable to combine different biosynthetic pathways *in situ*, by introducing SZs at the genomic level – *i.e.* applying CRISPR/Cas9 based genetic engineering. We are convinced that further research into this direction, like elucidating structures of SZ connected NRPS domains, eventually will bring up even more versatile artificial DDs as it is already suggested by NRPS-21 – 23 constructed with an synthetic (GS)_x_ C-A linker region.

## Acknowledgements

This work was supported by the LOEWE research cluster MegaSyn funded by the state of Hesse and an ERC advanced grant (grant agreement number 835108).

