## Supplementary material for "Synthetic Zippers as an Enabling Tool for Engineering of Non-Ribosomal Peptide Synthetases": Materials and Methods, Supplementary Tables and Figures

### Supplementary Information - Table of Contents

### 1 Material and methods

#### 1.1 Cultivation of strains

All *E. coli* DH10B::*mtaA*, *Xenorhabdus szentirmaii, Xenorhabdus nematophila* and *Photorhabdus luminescens* cells were cultivated in liquid or on solid LB-medium (pH 7.5, 10 g/L tryptone, 5 g/L yeast extract and 5 g/L NaCl). Solid media contained 1% (w/v) agar. Kanamycin (50 µg/ml), chloramphenicol (34 µg/ml) and spectinomycin (50 µg/ml) were used as selection markers. All *E. coli* cells cultures were cultivated at 37 °C and at 22 °C for peptide production purposes. *Xenorhabdus* and *Photorhabdus* strains were grown at 30 °C.

#### 1.2 Cloning of biosynthetic gene clusters

Genomic DNA of selected *Xenorhabdus* and *Photorhabdus* strains were isolated using the Qiagen Gentra Puregene Yeast/Bact Kit. All PCRs were performed with oligonucleotides obtained from Eurofins Genomics (Supplementary Table 4). NRPS fragments for Hot Fusion cloning (*1*) were amplified with primers coding for the respective homology arms (20-30 bp) in a two-step PCR program. The coding sequences for the SYNZIPs were also attached upstream or downstream to the NRPS genes by PCR. In the following, the cloning procedure for the basic vectors is explained. pJW61/62 was obtained by the following steps: First, the SYNZIP17/18 coding sequences (pENTR-SYNZIP17/18 (*2*) were a gift from Amy Keating, Addgene plasmids #80671/80672; RRID:Addgene_80671/80672) were inserted into the plasmids pCOLA_ara/tacI and pCK_0402 by oligonucleotides KB-pACYC-FW/RV or KB-pCOLA-FW/RV in two-step polymerase chain reactions (PCRs) combined with Hot Fusion Cloning (*1*). Second, these plasmids were linearized by single-step PCRs with the help of the oligonucleotides KB-pCOLA-II-FW/RV or KB-pACYC-II-FW/RV, which further allowed us to introduce NRPS fragments by Hot Fusion cloning. Therefore, the respective NRPS coding sequences were amplified again in two-step PCRs, using oligonucleotides with additional coding regions for homology arms (20-30 bp). pJW63/64 coding for subunits of the XtpS without attached SYNZIPs were generated by amplifying pJW61/62 with a single phosphorylated [phos.] oligonucleotide pair excluding the SYNZIP coding region followed by T4 DNA ligation (following Thermo Fisher manufacturers' instructions). The control plasmids pCOLA_ara_xtpS/gxpS_tacI_JW coding for the native single protein xtpS/ gxpS were created by Hot Fusion Cloning. Therefore, the plasmid pCOLA_ara/tacI was linearized by PCR using the oligonucleotides AL-XtpS-2-1 and AD64 and the insert *xtpS* was PCR amplified with the oligonucleotides jw0136_FW and jw0137_RV.

The plasmids pJW101/102 coding for NRPSs with two attached SNYZIPs were created by Hot Fusion cloning. Before this final cloning step an pCOLA_ara/tacI plasmid carrying the SYNZIP18 sequence downstream the $P_{BAD}$ promoter was linearized. This linearization step by PCR was done twice and allowed us to incorporate the SYNZIP1 coding region (na28_FW, na29_FW) upstream the stop codon (pQLinkHD-SYNZIP1 was a gift from Amy Keating (*2*), Addgene plasmid #80647; RRID:Addgene_80647).

The starting point for plasmid pJW103/106 was vector pCDF_ara/tacI. This vector was generated by digesting the plasmids pCOLA_ara/tacI and pCDFDuet™-1 (Novagen) with the enzymes XbaI and NdeI. The fragment of pCDFDuet™-1 carrying the pCloDF13 replicon and streptomycin/spectinomycin resistance marker was then T4 ligated with the compatible pCOLA_ara/tacI fragment. Then, this plasmid pCDF_ara/tacI was linearized in two cycles including in parallel the sequence of SYNZIP2 (na32_RV, na33_RV) (pQLinkHD-SYNZIP2 was a gift from Amy Keating (*2*), Addgene plasmid #80658; RRID:Addgene_80658) downstream the $P_{BAD}$ promoter, followed by the incorporation of the respective NRPS coding regions by Hot Fusion Cloning.

The plasmid pCOLA_ara_ gxpS_tacI_JW was generated in two Hot Fusion Cloning steps. First, the pCOLA_ara/tacI was linearized by PCR using the primers JW_tacI_PstI_FW2 and jw0064_RV and second the first part of *gxpS* was amplified using the oligonucleotides jw0124_FW/jw0160_RV. This intermediate plasmid was then opened with PstI and the second *gxpS* part, amplified with jw0151_FW/ jw0161_RV by PCR, was then integrated into the cleaving site by Hot Fusion Cloning.

In all PCRs the S7 Fusion High-Fidelity DNA Polymerase (Mobidiag) was used according to the manufacturers' instructions. The amplified DNA was purified with the Invisorb Fragment CleanUp or MSB Spin PCRapace Kits (stratec molecular). The basic cloning of all new generated plasmids (Supplementary Table 3) was performed in *E. coli* DH10B. Each NRPS (subunit) was under the control of a $P_{BAD}$ promotor. Plasmid isolation from *E. coli* was achieved with the Invisorb Spin Plasmid Mini Two Kit (stratec molecular). Restriction enzyme digests and the partial sequencing of essential plasmid regions, especially upstream and downstream of the NRPS genes, where the SYNZIP coding sequences were located, confirmed the correct plasmid construction.

#### 1.3 Heterologous expression of NRPS templates and LC-MS analysis

Constructed plasmids were transformed into *E. coli* DH10B::*mtaA*. Cells were grown overnight in LB medium containing the necessary antibiotics (50 µg/ml kanamycin, 34 µg/ml chloramphenicol, 50 µg/ml spectinomycin). 100 µl of an overnight culture were used for inoculation of 10 ml LB-cultures supplemented with the respective antibiotics as selection markers and additionally containing 0.002 mg/ml L-arabinose and 2 % (v/v) XAD-16. After incubation for 72 h at 22 °C the XAD-16 was harvested. One culture volume methanol was added and incubated for 60 min at 22 °C. The organic phase was filtrated and a sample was taken of the cleared extract. After centrifugation (17,000 x *g*, 20 min) the methanol extracts were used for LC-MS analysis. All measurements were performed by using an Ultimate 3000 LC system (Dionex) with an ACQUITY UPLC BEH C18 column (130 Å, 2.1 x 50 mm, 1.7 µm particle size; Waters) at a flow rate of 0.4 ml $min^{-1}$ using acetonitrile (ACN) and water containing 0.1% formic acid (v/v) in a gradient ranging from 5-95% of ACN over 16 min (40 °C) coupled to an AmaZonX (Bruker) electron spray ionization mass spectrometer. The BPC spectra were recorded in positive ion mode with the range from 100-1200 *m/z* and ultraviolet (UV) at 200-600 nm. The software Compass DataAnalysis 4.3 (Bruker) was used to evaluate the measurements.

#### 1.4 Peptide quantification

The absolute production titers of selected peptides were calculated with calibration curves based on pure synthetic **1** (for quantification of **1**, **10**, **11**, **22**, **23**, **27**), **2** (for quantification of **2-5**, **7-9, 25**, **26**), **6** (for quantification of **6** and **12**), **13** (for quantification of **13** and **14**), **15** (for quantification of **15** and **16**), **17**, **19** (for quantification of **18-21**), **24** (for quantification of **24**, **30**, **31**, **32**), **28** (for quantification of **28**, **29**) and **35** (for quantification of **33/34** and **35/36**). Therefore, the pure compounds were prepared at different concentrations (100, 50, 25, 12.5, 6.25, 3.125, 1.56, 0.78, 0.39, 0.195 and 0.0195 µg/mL) and measured by LC-MS using HPLC/MS measurements as described above. The peak area for each compound at different concentrations was calculated using Compass Data Analysis and used for the calculation of a standard curve passing through the origin. Triplicates of all *in vivo* experiments were measured. The pure peptide standards **1**, **2**, **6, 13**, **17** and **35** were synthesized in-house (*3, 4*) and the further pure synthetic **15**, **19**, **24** and **28** were produced by Synpeptide.

#### 1.5 Chemical synthesis

Chemical synthesis of all peptides was performed as described previously (*4*).

#### 2 Supplementary Tables

**Table S1.** ESI-MS data of all produced peptides.

| Peptide (#) | theoretical mass-to-charge ratio (*m/z*) $[M+H]^+$ | Molecular formula | Reference |
|---|---|---|---|
| **1** | 410.29 | $C_{21}H_{38}N_4O_4$ | (*5*) |
| **2** | 586.40 | $C_{32}H_{51}O_5N_5$ | (*6*) |
| **3** | 600.41 | $C_{33}H_{53}O_5N_5$ | (*6*) |
| **4** | 552.41 | $C_{29}H_{53}O_5N_5$ | (*6*) |
| **5** | 566.43 | $C_{30}H_{55}O_5N_5$ | (*6*) |
| **6** | 556.35 | $C_{27}H_{49}N_5O_5S$ | - |
| **7** | 556.41 | $C_{28}H_{53}N_5O_6$ | - |
| **8** | 570.42 | $C_{29}H_{55}N_5O_6$ | - |
| **9** | 584.44 | $C_{30}H_{57}N_5O_6$ | - |
| **10** | 457.34 | $C_{23}H_{44}N_4O_5$ | - |
| **11** | 471.35 | $C_{24}H_{46}N_4O_5$ | - |
| **12** | 556.35 | $C_{27}H_{49}N_5O_5S$ | - |
| **13** | 589.33 | $C_{29}H_{44}N_6O_7$ | - |
| **14** | 555.35 | $C_{26}H_{46}N_6O_7$ | - |
| **15** | 634.38 | $C_{32}H_{51}N_5O_8$ | (*4*) |
| **16** | 600.40 | $C_{29}H_{53}N_5O_8$ | (*4*) |
| **17** | 643.43 | $C_{33}H_{54}N_8O_5$ | - |
| **18** | 830.54 | $C_{43}H_{71}N_7O_9$ | - |
| **19** | 844.55 | $C_{44}H_{73}N_7O_9$ | - |
| **20** | 858.57 | $C_{45}H_{75}N_7O_9$ | - |
| **21** | 810.57 | $C_{41}H_{75}N_7O_9$ | - |
| **22** | 459,30 | $C_{25}H_{38}N_4O_4$ | - |
| **23** | 425,31 | $C_{22}H_{40}N_4O_4$ | - |
| **24** | 778,45 | $C_{41}H_{59}N_7O_8$ | - |
| **25** | 538.40 | $C_{28}H_{51}N_5O_5$ | - |
| **26** | 552.41 | $C_{29}H_{53}N_5O_5$ | - |
| **27** | 425.31 | $C_{22}H_{40}N_4O_4$ | - |
| **28** | 826.45 | $C_{45}H_{59}N_7O_8$ | - |
| **29** | 840.47 | $C_{46}H_{61}N_7O_8$ | - |
| **30** | 792.47 | $C_{42}H_{61}N_7O_8$ | - |
| **31** | 806.48 | $C_{43}H_{63}N_7O_8$ | - |
| **32** | 792.47 | $C_{42}H_{61}N_7O_8$ | - |
| 33 | 358.27 | $C_{18}H_{35}N_3O_4$ | - |
| 34 | 358.27 | $C_{18}H_{35}N_3O_4$ | - |
| 35 | 392.25 | $C_{21}H_{33}N_3O_4$ | - |
| 36 | 392.25 | $C_{21}H_{33}N_3O_4$ | - |
| 37 | 314.27 | $C_{18}H_{35}NO_3$ | - |
| 38 | 328.29 | $C_{19}H_{37}NO_3$ | - |
| 39 | 342.20 | $C_{20}H_{39}NO_3$ | - |
| 40 | 455.38 | $C_{26}H_{50}N_2O_4$ | - |
| 41 | 510.39 | $C_{28}H_{51}N_3O_5$ | (*7*) |
| 42 | 496.37 | $C_{27}H_{49}N_3O_5$ | (*7*) |
| 43 | 500.20 | $C_{26}H_{49}N_3O_6$ | - |

**Table S2.** Strains used in this work.

| Strain | Genotype / NRPS | Reference |
|---|---|---|
| *E. coli* DH10B | F_mcrA (*mrr-hsd*RMS-*mcr*BC), 80*lac*ZΔ, M15, Δ*lac*X74 *rec*A1 *end*A1 *ara*D 139Δ(*ara*, *leu*)7697 *gal*U *gal*K λ *rps*L (*Strr*) *nup*G / - | (*8*) |
| *E. coli* DH10B::*mtaA* | DH10B with *mtaA* from pCK_*mtaA*Δ*entD* / - | (*9*) |
| *P. luminescens* TTO1 | - / *gxpS* (*6*) | DSMZ |
| *X. bovienii* SS-2004 | - / *garS*/ *xfpS (7)* | (*10*) |
| *X. nematophila* ATCC 19061 | - / *xtpS (5)* | ATCC |
| *X. budapestensis* DSM 16342 | - / *bicA* (*11*) | DSMZ |
| *X. miraniensis* DSM 17902 | - / *ambS* (*9*) | DSMZ |
| *X. szentirmaii* DSM16338 | - / *szeS* | DSMZ |
| *X. indica* DSM 17382 | - / *xldS* (*9*) | DSMZ |
| *B. licheniformis* ATCC 10716 | - / *bacA (12)* | M. A. Marahiel / ATCC |
| *B. subitlis* MR 168 | - / *srfA (13)* | ATCC |

**Table S3.** Plasmids used in this work.

| Plasmids | Genotype | Reference |
|---|---|---|
| pFF1_22A_szeS_gxpS | ori 2µ, kanMX4, ori ColA, kan$^R$, $P_{BAD}$ *szeS*_Ft$A_1T_1C/E_2A_2T_2C_3$-*gxpS*_$A_3T_3C/E_4A_4T_4C/E_5A_5T_5TE$, Ypet-Flag | (*4*) |
| pFF1_NRPS_6 | ori 2µ, kanMX4, *araC-P*$_{BAD}$, ori ColA, Ypet-Flag, kan$^R$, *bacA*-A1T1CyA2T2C3A3T3CD$_{sub4}$_*sfrA-BC*-C$_{Asub6}$A6T6E6C7A7T7TE | (*3*) |
| pCOLA_ara/tacI | ori ColA, kan$^R$, *araC-P*$_{BAD}$ and *tac*I | (*14*) |
| pCK_0402 | ori p15A, cm$^R$, *araC-P*$_{BAD}$ and *tac*I-*araE* | unpublished |
| pCDF_ara/tacI | ori CloDF13, spec$^R$, *araC-P*$_{BAD}$ and *tac*I | this study |
| pCOLA_ara_xtpS_tacI_JW | ori ColA, kan$^R$, *araC-P*$_{BAD}$ *xtpS* and *tac*I | this study |
| pCOLA_ara_gxpS_tacI_JW | ori ColA, kan$^R$, *araC-P*$_{BAD}$ *gxpS* and *tac*I | this study |
| pJW61 | ori p15A, cm$^R$, *araC-P*$_{BAD}$ *xtpS*_$A_1T_1C/E_2A_2T_2C_3$-SYNZIP17 and *tac*I-*araE* | this study |
| pJW62 | ori ColA, kan$^R$, *araC-P*$_{BAD}$ SYNZIP18-*xtpS*_$A_3T_3C/E_4A_4T_4TE$ and *tac*I | this study |
| pJW63 | ori p15A, cm$^R$, *araC-P*$_{BAD}$ *xtpS*_$A_1T_1C/E_2A_2T_2C_3$ and *tac*I-*araE* | this study |
| pJW64 | ori ColA, kan$^R$, *araC-P*$_{BAD}$ *xtpS*_$A_3T_3C/E_4A_4T_4TE$, *tac*I | this study |
| pJW75 | ori p15A, cm$^R$, *araC-P*$_{BAD}$ *gxpS*_$A_1T_1C/E_2A_2T_2C_3$-SYNZIP17 and *tac*I-*araE* | this study |
| pJW76 | ori ColA, kan$^R$, *araC-P*$_{BAD}$ SYNZIP18-*gxpS*_$A_3T_3C/E_4A_4T_4C/E_5A_5T_5TE$ and *tac*I | this study |
| pJW77 | ori p15A, cm$^R$, *araC-P*$_{BAD}$ *bicA*_$A_1T_1C/E_2A_2T_2C_3$-SYNZIP17 and *tac*I-*araE* | this study |
| pJW91 | ori p15A, cm$^R$, *araC-P*$_{BAD}$ *ambS*_ $A_1T_1C/E_2A_2T_2C_3$-SYNZIP17 and *tac*I-*araE* | this study |
| pJW92 | ori p15A, cm$^R$, *araC-P*$_{BAD}$ *szeS*_Ft$A_1T_1C/E_2A_2T_2C_3$-SYNZIP17 and *tac*I-*araE* | this study |
| pJW93 | ori p15A, cm$^R$, *araC-P*$_{BAD}$ *xldS*_$C_1A_1T_1C/E_2A_2T_2C_3$-SYNZIP17 and *tac*I-*araE* | this study |
| pJW100 | ori p15A, cm$^R$, *araC-P*$_{BAD}$ *xtpS*_$A_1T_1C/E_2$-$(GS)_2$-SYNZIP17 and *tac*I-*araE* | this study |
| pJW102 | ori ColA, kan$^R$, *araC-P*$_{BAD}$ SYNZIP18-*xtpS*_$A_2T_2C_3$-$(GS)_2$-SYNZIP1 and *tac*I | this study |
| pJW103 | ori CloDF13, spec$^R$, *araC-P*$_{BAD}$ *xtpS*_SYNZIP2-$A_3T_3C/E_4A_4T_4TE$ and *tac*I | this study |
| pJW114 | ori p15A, cm$^R$, *araC-P*$_{BAD}$ *bacA*_A1T1CyA2T2C3-SYNZIP17 and *tac*I-*araE* | this study |
| pJW116 | ori ColA, kan$^R$, *araC-P*$_{BAD}$ SYNZIP18-*bacA*_A3T3C$_{Dsub}$4-*sfrA-BC*_C$_{Asub}$6A6T6E6C7A7T7TE and *tac*I | this study |
| pJW118 | ori p15A, cm$^R$, *araC-P*$_{BAD}$ *bacA*_A1T1CyA2T2C3A3-SYNZIP17 and *tac*I-*araE* | this study |
| pJW120 | ori ColA, kan$^R$, *araC-P*$_{BAD}$ SYNZIP18-*bacA*_T3C$_{Dsub}$4-*sfrA-BC*_C$_{Asub}$6A6T6E6C7A7T7TE and *tac*I | this study |
| pJW122 | ori p15A, cm$^R$, *araC-P*$_{BAD}$ *bacA*_A1T1CyA2T2C3A3T3-SYNZIP17 and *tac*I-*araE* | this study |
| pJW124 | ori ColA, kan$^R$, *araC-P*$_{BAD}$ SYNZIP18-*bacA*_C$_{Dsub}$4-*sfrA-BC*_C$_{Asub}$6A6T6E6C7A7T7TE and *tac*I | this study |
| pJW126 | ori p15A, cm$^R$, *araC-P*$_{BAD}$ *bacA*_A1T1CyA2T2C3A3T3 C$_{Dsub}$4-SYNZIP17 and *tac*I-*araE* | this study |
| pJW128 | ori ColA, kan$^R$, *araC-P*$_{BAD}$ SYNZIP18-*sfrA-BC*_C$_{Asub}$6A6T6E6C7A7T7TE and *tac*I | this study |
| pJW141 | ori p15A, cm$^R$, *araC-P*$_{BAD}$ *xldS*_$C_1$-SYNZIP17 and *tac*I-*araE* | this study |
| pNA1 | ori p15A, cm$^R$, *araC-P*$_{BAD}$ *xtpS*_ $A_1T_1C/E_2A_2T_2C_3$-SYNZIP19 and *tac*I-*araE* | this study |
| pNA2 | ori p15A, cm$^R$, *araC-P*$_{BAD}$ *xtpS*_ $A_1T_1C/E_2A_2T_2$-SYNZIP17 und *tac*I-*araE* | this study |
| pNA3 | ori ColA, kan$^R$, *araC-P*$_{BAD}$ SYNZIP18-*xtpS*_ $C_3A_3T_3C/E_4A_4T_4TE$ und *tac*I | this study |
| pNA4 | ori p15A, cm$^R$, *araC-P*$_{BAD}$ *xtpS*_ $A_1T_1C/E_2A_2$-SYNZIP17 und *tac*I-*araE* | this study |

| | | |
|---|---|---|
| pNA5 | ori ColA, kan$^R$, *araC-P$_{BAD}$* SYNZIP18-*xtpS*_ $T_2C_3A_3T_3C/E_4A_4T_4$TE und *tac*I | this study |
| pNA8 | ori p15A, cm$^R$, *araC-P$_{BAD}$* *xtpS*_$A_1T_1C/E_2A_2T_2C_3$-$(GS)_5$-SYNZIP17 and *tac*I-*araE* | this study |
| pNA9 | ori p15A, cm$^R$, *araC-P$_{BAD}$* *xtpS*_$A_1T_1C/E_2A_2T_2C_3$-$(GS)_4$-SYNZIP17 and *tac*I-*araE* | this study |
| pNA10 | ori p15A, cm$^R$, *araC-P$_{BAD}$* *xtpS*_$A_1T_1C/E_2A_2T_2C_3$-$(GS)_2$-SYNZIP17 and *tac*I-*araE* | this study |
| pNA15 | ori ColA, kan$^R$, *araC-P$_{BAD}$* SYNZIP18-*xtpS*_$T_2C_3A_3$-SYNZIP1 and *tac*I | this study |
| pNA16 | ori CloDF13, spec$^R$, *araC-P$_{BAD}$* SYNZIP2-*xtpS*_$T_3C/E_4A_4T_4$TE and *tac*I | this study |
| pNA17 | ori ColA, kan$^R$, *araC-P$_{BAD}$* SYNZIP18-*xtpS*_$C_3A_3T_4$-SYNZIP1 and *tac*I | this study |
| pNA18 | ori CloDF13, spec$^R$, *araC-P$_{BAD}$* SYNZIP2-*xtpS*_$C/E_4A_4T_4$TE and *tac*I | this study |
| pNA26 | ori p15A, cm$^R$, *araC-P$_{BAD}$* *gxpS*_$A_1T_1C/E_2A_2$-SYNZIP17 and *tac*I-*araE* | this study |
| pNA27 | ori ColA, kan$^R$, *araC-P$_{BAD}$* SYNZIP18-*gxpS*_$T_2C_3A_3$-SYNZIP1 and *tac*I | this study |
| pNA28 | ori CloDF13, spec$^R$, *araC-P$_{BAD}$* SYNZIP2-*gxpS*_$T_3C/E_4A_4T_4$ $C/E_5A_5T_5$TE and *tac*I | this study |
| pNA29 | ori p15A, cm$^R$, *araC-P$_{BAD}$* *sze*S_ $C_1A_1T_1C/E_2A_2$-SYNZIP17 and *tac*I-*araE* | this study |
| pNA30 | ori ColA, kan$^R$, *araC-P$_{BAD}$* SYNZIP18-*szeS*_$T_2C_3A_3$-SYNZIP1 and *tac*I | this study |
| pNA31 | ori CloDF13, spec$^R$, *araC-P$_{BAD}$* SYNZIP2-*szeS*_$T_3C/E_4A_4T_4$ $C/E_5A_5T_5C_6A_6T_6$TE and *tac*I | this study |
| pNA35 | ori ColA, kan$^R$, *araC-P$_{BAD}$* SYNZIP18-*garS*_$T_2C_3A_3$-SYNZIP1 and *tac*I | this study |
| pNA59 | ori p15A, cm$^R$, *araC-P$_{BAD}$* *xfpS*_$C_1A_1$-SYNZIP17 and *tac*I-*araE* | this study |
| pNA60 | ori ColA, kan$^R$, *araC-P$_{BAD}$* SYNZIP18-*xfpS*_$T_1E_1C_2A_2T_2C_3A_3T_3$TE and *tac*I | this study |

**Table S4.** Oligonucleotides used in this work. Correlations of plasmids to figures from the main text and supplementary informationt are represented in brackets.

| Plasmids | Oligo-nucleotide | Sequence (5´→3´; overlapping ends) | Template |
|---|---|---|---|
| pJW61 (NRPS-1, NRPS-2, NRPS-8, NRPS-9) | KB-pACYC-FW | GAACAGTTAAAACAGAAGCGTGAACAATTAAAGCAAAAGATCGCCAATCTGCGTAA GGAGATCGAAGCCTACAAGTGACAATTAATCATCGGCTCG | pCK_0402 |
| | KB-pACYC-RV | TTCACGCTTCTGTTTTAACTGTTCGATGCGATTACGCAATTCAGCCTTTTTCGATTTT AATTCCTCCTTCTCGTTCATGGAATTCCTCCTGTTAGC | pCK_0402 |
| | KB-pACYC-II-FW | AACGAGAAGGAGGAATTAAAATCG | - |
| | KB-pACYC-II-RV | CATGGAATTCCTCCTGTTAGC | - |
| | KB-P1-FW | TGGGCTAACAGGAGGAATTCCATGAAAGATAGCATGGCTAAAAAGGG | *X. nematophila* ATCC 19061 |
| | KB-P1-RV | CGATTTTAATTCCTCCTTCTCGTTCCAGGTTTTTAACAACAATGTGC | *X. nematophila* ATCC 19061 |
| pJW62 (NRPS-1, NRPS-3, NRPS-7, NRPS-12, NRPS-17, NRPS-18, NRPS-19) | KB-pCOLA-FW | CATTGACAAAGAGCTGCGTGCCAACGAAAACGAACTTCGCGCCCTTGATAACGAGC TGACTGCAGCTATCTCATGACAATTAATCATCGGCTCG | pCOLA_ara/tacI |
| | KB-pCOLA-RV | TTGGCACGCAGCTCTTTGTCAATGGCATTTAACTCGCGGTCCAAGGCTTTCAGTTCA CGCTCTTCAGCATAGAACATGGAATTCCTCCTGTTAGC | pCOLA_ara/tacI |
| | KB-pCOLA-II-FW | TGACAATTAATCATCGGCTCG | - |
| | KB-pCOLA-II-RV | TGAGATAGCTGCAGTCAGCTCG | - |
| | KB-P2-FW | AACGAGCTGACTGCAGCTATCTCATTATGTATTCATCAACTTTTTGAACAGC | *X. nematophila* ATCC 19061 |
| | KB-P2-RV | ATACGAGCCGATGATTAATTGTCACAGCGCCTCCACTTCG | *X. nematophila* ATCC 19061 |
| pJW63 (NRPS-3, NRPS-4) | jw0061_FW | [phos.] TGACAATTAATCATCGGCTCG | pJW61 |
| | jw0062_RV | CCAGGTTTTTAACAACAATGTGC | pJW61 |
| pJW64 (NRPS-2, NRPS-4) | jw0063_FW | [phos.] TTATGTATTCATCAACTTTTTGAACAGC | pJW62 |
| | jw0064_RV | CATGGAATTCCTCCTGTTAGCC | pJW62 |
| pJW75 (NRPS-5, NRPS-7, NRPS-10) | KB-pACYC-II-FW | AACGAGAAGGAGGAATTAAAATCG | pJW61 |
| | KB-pACYC-II-RV | CATGGAATTCCTCCTGTTAGC | pJW61 |
| | jw0124_FW | GGGCTAACAGGAGGAATTCCATGAAAGATAGCATGGCTAAAAAGGAAATTATC | *P. luminescens* TTO1 |
| | jw0125_RV | TCGATTTTAATTCCTCCTTCTCGTTCCAATTTTCCAGTAATAACTCCCG | *P. luminescens* TTO1 |
| pJW76 (NRPS-5, NRPS-8, NRPS-11, NRPS-13, NRPS-14, NRPS-15, NRPS-16) | KB-pCOLA-II-FW | TGACAATTAATCATCGGCTCG | pJW62 |
| | KB-pCOLA-II-RV | TGAGATAGCTGCAGTCAGCTCG | pJW62 |
| | jw0172_FW | GGCTAACAGGAGGAATTCCATGTTCTATGCTGAAGAGCGTGAAC | *P. luminescens* TTO1 |
| | jw0127_RV | CGAGCCGATGATTAATTGTCACAGCGCCTCCGCTTC | *P. luminescens* TTO1 |
| pJW114 (NRPS-6, NRPS-11, NRPS-12) | KB-pACYC-II-FW | AACGAGAAGGAGGAATTAAAATCG | pJW61 |
| | KB-pACYC-II-RV | CATGGAATTCCTCCTGTTAGC | pJW61 |
| | jw208_FW | GCTAACAGGAGGAATTCCATGGTTGCTAAACATTCATTAGAAAATGGG | pFF1_NRPS_6 (*3*) |
| | jw209_RV | CGATTTTAATTCCTCCTTCTCGTTCTTTGTATGGTTAAAGGACTCTAAAAGTGTC | pFF1_NRPS_6 (*3*) |
| pJW116 (NRPS-6, NRPS-9, NRPS-10, NRPS-43) | KB-pCOLA-II-FW | TGACAATTAATCATCGGCTCG | pJW62 |
| | KB-pCOLA-II-RV | TGAGATAGCTGCAGTCAGCTCG | pJW62 |
| | jw0211_FW | CGAGCTGACTGCAGCTATCTCAAAAGCAATCCACCAGCTGTTT | pFF1_NRPS_6 (*3*) |
| | jw0212_RV | CGAGCCGATGATTAATTGTCATGAAACCGTTACGGTTTGTGTATTA | pFF1_NRPS_6 (*3*) |
| pJW77 (NRPS-15) | KB-pACYC-II-FW | AACGAGAAGGAGGAATTAAAATCG | pJW61 |
| | KB-pACYC-II-RV | CATGGAATTCCTCCTGTTAGC | pJW61 |
| | jw0128_FW | GGGCTAACAGGAGGAATTCCATGAAAGATAACATTGCTACAGTGGCAAATAG | *X. budapestensis* DSM 16342 |
| | jw0129_RV | CGATTTTAATTCCTCCTTCTCGTTCCAAGTTTTCAGCAACAACTGG | *X. budapestensis* DSM 16342 |
| pJW91 (NRPS-13, NRPS-17) | KB-pACYC-II-FW | AACGAGAAGGAGGAATTAAAATCG | pJW61 |
| | KB-pACYC-II-RV | CATGGAATTCCTCCTGTTAGC | pJW61 |
| | jw0162_FW | GCTAACAGGAGGAATTCCATGAAAAATGATAAGGTGATGACTCTG | *X. miraniensis* DSM 17902 |
| | jw0163_RV | TCGATTTTAATTCCTCCTTCTCGTTCCACGTTTCCAGCAATAACC | *X. miraniensis* DSM 17902 |
| pJW92 (NRPS-14, NRPS-19) | KB-pACYC-II-FW | AACGAGAAGGAGGAATTAAAATCG | pJW61 |
| | KB-pACYC-II-RV | CATGGAATTCCTCCTGTTAGC | pJW61 |
| | jw0164_FW | GCTAACAGGAGGAATTCCATGAAAGGTAGTATTGCTAAAAAGGGAG | *X. szentirmaii* DSM16338 |
| | jw0165_RV | TCGATTTTAATTCCTCCTTCTCGTTCCAGCTTTCCAGCAATAACC | *X. szentirmaii* DSM16338 |
| pJW93 (NRPS-16, NRPS-18) | KB-pACYC-II-FW | AACGAGAAGGAGGAATTAAAATCG | pJW61 |
| | KB-pACYC-II-RV | CATGGAATTCCTCCTGTTAGC | pJW61 |

| | | | |
|---|---|---|---|
| | jw0166_FW | GCTAACAGGAGGAATTCCATGAAACTTTGGAACTATAAAATGAATATGAC | *X. indica* DSM 17382 |
| | jw0167_RV | TCGATTTTAATTCCTCCTTCTCGTTGAAATCCACCAACAGTTGTTGAC | *X. indica* DSM 17382 |
| pJW100 (NRPS-26) | KB-pACYC-II-FW | AACGAGAAGGAGGAATTAAAATCG | pJW61 |
| | KB-pACYC-II-RV | CATGGAATTCCTCCTGTTAGC | pJW61 |
| | KB-P1-FW | TGGGCTAACAGGAGGAATTCCATGAAAGATAGCATGGCTAAAAAGGG | *X. nematophila* ATCC 19061 |
| | jw0179_RV | CGATTTTAATTCCTCCTTCTCGTTGGATCCAGACCCCCAGGTTTTCAGCAATAACGTG | *X. nematophila* ATCC 19061 |
| pJW102 (NRPS-26) | na29_FW | AACCTGGTTGCGCAGCTCGAAAACGAAGTTGCGTCTCTGGAAAATGAGAACGAAACCCTGAAGAAAAAGAACCTGCACAAAAAAGACCTGATCGCGTAC | pJW101 |
| | KB-pCOLA-II-RV | TGAGATAGCTGCAGTCAGCTCG | pJW101 |
| | jw0180_FW | AACGAGCTGACTGCAGCTATCTCACTGTGTATCCATCAGTTAATTGAACAACAG | *X. nematophila* ATCC 19061 |
| | jw0182_RV | CGTTTTCGAGCTGCGCAACCAGGTTGGATCCAGACCCCCAGGTTTTTAACAACAATGTGCG | *X. nematophila* ATCC 19061 |
| pJW103 (NRPS-26) | na34_FW | TGACAATTAATCATCGGCTCG | pCDF_ara/tacI |
| | na32_RV | GTTCTGTTCATCACGTTCCAGCTGCAGGTTGTCTTTTTTCAGACGTGCGATTTTCTTACGCAGATACGCGTTACGCGCCATGGAATTCCTCCTGTTAGCC | pCDF_ara/tacI |
| | na33_RV | CTGTTCGTGAGACGCAACTTCGTTTTCGAGACGCGCGATTTCGTCACGCAGGTTCGCGATGATTTTTTCCAGGTTCTGTTCATCACGTTCCAGC | pCDF_ara/tacI |
| | na84_RV | CTGTTCGTGAGACGCAACTTC | - |
| | jw0183_FW | AAACGAAGTTGCGTCTCACGAACAGTTATGTATTCATCAACTTTTTGAACAGC | *X. nematophila* ATCC 19061 |
| | jw0188_RV | GCCTAAACCAATACGCCGT | *X. nematophila* ATCC 19061 |
| | jw0189_FW | CGGCGTATTGGTTTAGGCCTGT | *X. nematophila* ATCC 19061 |
| | na07_RV | CGAGCCGATGATTAATTGTCACAGCGCCTCCACTTCG | *X. nematophila* ATCC 19061 |
| pJW118 (NRPS-44) | KB-pACYC-II-FW | AACGAGAAGGAGGAATTAAAATCG | pJW61 |
| | KB-pACYC-II-RV | CATGGAATTCCTCCTGTTAGC | pJW61 |
| | jw208_FW | GCTAACAGGAGGAATTCCATGGTTGCTAAACATTCATTAGAAAATGGG | pFF1_NRPS_6 (*3*) |
| | jw0214_RV | CGATTTTAATTCCTCCTTCTCGTTGTAGCGGCGATCCATTGT | pFF1_NRPS_6 (*3*) |
| pJW120 (NRPS-44) | KB-pCOLA-II-FW | TGACAATTAATCATCGGCTCG | pJW62 |
| | KB-pCOLA-II-RV | TGAGATAGCTGCAGTCAGCTCG | pJW62 |
| | jw0216_FW | CGAGCTGACTGCAGCTATCTCAGAAGCGCCGCGGG | pFF1_NRPS_6 (*3*) |
| | jw0212_RV | CGAGCCGATGATTAATTGTCATGAAACCGTTACGGTTTGTGTATTA | pFF1_NRPS_6 (*3*) |
| pJW122 (NRPS-45) | KB-pACYC-II-FW | AACGAGAAGGAGGAATTAAAATCG | pJW61 |
| | KB-pACYC-II-RV | CATGGAATTCCTCCTGTTAGC | pJW61 |
| | jw208_FW | GCTAACAGGAGGAATTCCATGGTTGCTAAACATTCATTAGAAAATGGG | pFF1_NRPS_6 (*3*) |
| | jw0218_RV | CGATTTTAATTCCTCCTTCTCGTTTCGGTAATATGGTTTTTCTTCGG | pFF1_NRPS_6 (*3*) |
| pJW124 (NRPS-45) | KB-pCOLA-II-FW | TGACAATTAATCATCGGCTCG | pJW62 |
| | KB-pCOLA-II-RV | TGAGATAGCTGCAGTCAGCTCG | pJW62 |
| | jw0220_FW | CGAGCTGACTGCAGCTATCTCA TTGTCTTCAGCGCAAAAAAGG | pFF1_NRPS_6 (*3*) |
| | jw0212_RV | CGAGCCGATGATTAATTGTCA TGAAACCGTTACGGTTTGTGTATTA | pFF1_NRPS_6 (*3*) |
| pJW126 (NRPS-46) | KB-pACYC-II-FW | AACGAGAAGGAGGAATTAAAATCG | pJW61 |
| | KB-pACYC-II-RV | CATGGAATTCCTCCTGTTAGC | pJW61 |
| | jw208_FW | GCTAACAGGAGGAATTCCATGGTTGCTAAACATTCATTAGAAAATGGG | pFF1_NRPS_6 (*3*) |
| | jw0222_RV | CGATTTTAATTCCTCCTTCTCGTTGGCATGGCTATTTTCCCATT | pFF1_NRPS_6 (*3*) |
| pJW128 (NRPS-46) | KB-pCOLA-II-FW | TGACAATTAATCATCGGCTCG | pJW62 |
| | KB-pCOLA-II-RV | TGAGATAGCTGCAGTCAGCTCG | pJW62 |
| | jw0224_FW | CGAGCTGACTGCAGCTATCTCACAAAAAGAACGGATGAAGGAGC | pFF1_NRPS_6 (*3*) |
| | jw0212_RV | CGAGCCGATGATTAATTGTCATGAAACCGTTACGGTTTGTGTATTA | pFF1_NRPS_6 (*3*) |
| pJW141 (NRPS-43) | KB-pACYC-II-FW | AACGAGAAGGAGGAATTAAAATCG | pJW61 |
| | KB-pACYC-II-RV | CATGGAATTCCTCCTGTTAGC | pJW61 |
| | jw0166_FW | GCTAACAGGAGGAATTCCATGAAACTTTGGAACTATAAAATGAATATGAC | *X. indica* DSM 17382 |
| | jw0254_RV | CGATTTTAATTCCTCCTTCTCGTTAAAATCTACCAATAGTTTCTGGCGC | *X. indica* DSM 17382 |
| pNA1 (NRPS-20) | na01_FW | CGTGAACAGCTGAAACAGAAACTGGCGGCTCTGCGTAACAAACTGGACGCGTACAAAAACCGTCTG TGACAATTAATCATCGGCTCG | pCK_0402 |
| | na02_FW | AACGAACTGGAATCTCTGGAGAACAAAAAAGAAGAACTGAAGAACCGTAACGAAGAGCTGAAGCAGAAA CGTGAACAGCTGAAACAGAAAC | pCK_0402 |
| | jw0064_RV | CATGGAATTCCTCCTGTTAGCC | pCK_0402 |
| | na03_FW | TGGGCTAACAGGAGGAATTCCATGAAAGATAGCATGGCTAAAAAGGG | *X. nematophila* ATCC 19061 |

| | | | |
|---|---|---|---|
| | na04_RV | CTCCAGAGATTCCAGTTCGTTCCAGGTTTTTAACAACAATGTGC | *X. nematophila* ATCC 19061 |
| pNA2 (NRPS-24, NRPS-27) | KB-pACYC-II-FW | AACGAGAAGGAGGAATTAAAATCG | pJW61 |
| | KB-pACYC-II-RV | CATGGAATTCCTCCTGTTAGC | pJW61 |
| | na03_FW | TGGGCTAACAGGAGGAATTCCATGAAAGATAGCATGGCTAAAAAGGG | *X. nematophila* ATCC 19061 |
| | na05_RV | CGATTTTAATTCCTCCTTCTCGTTAACACGATCACGGGATATTG | *X. nematophila* ATCC 19061 |
| pNA3 (NRPS-24) | KB-pCOLA-II-FW | TGACAATTAATCATCGGCTCG | pJW62 |
| | KB-pCOLA-II-RV | TGAGATAGCTGCAGTCAGCTCG | pJW62 |
| | na06 | AACGAGCTGACTGCAGCTATCTCATTGCCTTTATCGTTTGGTCAAC | *X. nematophila* ATCC 19061 |
| | na07 | CGAGCCGATGATTAATTGTCACAGCGCCTCCACTTCG | *X. nematophila* ATCC 19061 |
| pNA4 (NRPS-25, NRPS-28, NRPS-29, NRPS-30, NRPS-31, NRPS-32, NRPS-33, NRPS-42) | KB-pACYC-II-FW | AACGAGAAGGAGGAATTAAAATCG | pJW61 |
| | KB-pACYC-II-RV | CATGGAATTCCTCCTGTTAGC | pJW61 |
| | na03_FW | TGGGCTAACAGGAGGAATTCCATGAAAGATAGCATGGCTAAAAAGGG | *X. nematophila* ATCC 19061 |
| | na13_RV | CGATTTTAATTCCTCCTTCTCGTTATAAATCTGGCGGGCGAA | *X. nematophila* ATCC 19061 |
| pNA5 (NRPS-25, NRPS-48) | KB-pCOLA-II-FW | TGACAATTAATCATCGGCTCG | pJW62 |
| | KB-pCOLA-II-RV | TGAGATAGCTGCAGTCAGCTCG | pJW62 |
| | na14_FW | AACGAGCTGACTGCAGCTATCTCAGTTGCGCCACAAGGAGAA | *X. nematophila* ATCC 19061 |
| | na07_RV | CGAGCCGATGATTAATTGTCACAGCGCCTCCACTTCG | *X. nematophila* ATCC 19061 |
| pNA8 (NRPS-23) | KB-pACYC-II-FW | AACGAGAAGGAGGAATTAAAATCG | pJW61 |
| | KB-pACYC-II-RV | CATGGAATTCCTCCTGTTAGC | pJW61 |
| | na3_FW | TGGGCTAACAGGAGGAATTCCATGAAAGATAGCATGGCTAAAAAGGG | *X. nematophila* ATCC 19061 |
| | na17_RV | CGATTTTAATTCCTCCTTCTCGTTTGATCCCGAACCTGAGCCGGATCCAGACCCCCAGGTTTTTAACAACAATGTGC | *X. nematophila* ATCC 19061 |
| pNA9 (NRPS-22) | KB-pACYC-II-FW | AACGAGAAGGAGGAATTAAAATCG | pJW61 |
| | KB-pACYC-II-RV | CATGGAATTCCTCCTGTTAGC | pJW61 |
| | na3_FW | TGGGCTAACAGGAGGAATTCCATGAAAGATAGCATGGCTAAAAAGGG | *X. nematophila* ATCC 19061 |
| | na19_RV | CGATTTTAATTCCTCCTTCTCGTTCGAACCTGAGCCGGATCCAGACCCCCAGGTTTTTAACAACAATGTGC | *X. nematophila* ATCC 19061 |
| pNA10 (NRPS-21) | KB-pACYC-II-FW | AACGAGAAGGAGGAATTAAAATCG | pJW61 |
| | KB-pACYC-II-RV | CATGGAATTCCTCCTGTTAGC | pJW61 |
| | na3_FW | TGGGCTAACAGGAGGAATTCCATGAAAGATAGCATGGCTAAAAAGGG | *X. nematophila* ATCC 19061 |
| | na20_RV | CGATTTTAATTCCTCCTTCTCGTTGGATCCAGACCCCCAGGTTTTTAACAACAATGTG | *X. nematophila* ATCC 19061 |
| pNA15 (NRPS-28, NRPS-29, NRPS-32, NRPS-37, NRPS-38, NRPS-40) | na29_FW | AACCTGGTTGCGCAGCTCGAAAACGAAGTTGCGTCTCTGGAAAATGAGAACGAAACCCTGAAGAAAAAGAACCTGCACAAAAAAGACCTGATCGCGTAC | pJW61 |
| | na30_RV | TGAGATAGCTGCAGTCAGCTCG | pJW61 |
| | na22_FW | GGCTAACAGGAGGAATTCCATGGTTGCGCCACAAGGAGAA | *X. nematophila* ATCC 19061 |
| | na41_RV | CGAGCCGATGATTAATTGTCAATAGACCTGCCGGGCAAAC | *X. nematophila* ATCC 19061 |
| pNA16 (NRPS-28, NRPS-30, NRPS-31, NRPS-35, NRPS-38) | na33_RV | CTGTTCGTGAGACGCAACTTCGTTTTCGAGACGCGCGATTTCGTCACGCAGGTTCGCGATGATTTTTTCCAGGTTCT GTTCATCACGTTCCAGC | pCDF_ara/tacI |
| | na34_FW | TGACAATTAATCATCGGCTCG | pCDF_ara/tacI |
| | na42_FW | TTGGGCTAACAGGAGGAATTCC ATGGCGGCTCCGCAGGG | *X. nematophila* ATCC 19061 |
| | na7_RV | CGAGCCGATGATTAATTGTCA CAGCGCCTCCACTTCG | *X. nematophila* ATCC 19061 |
| pNA17 (NRPS-27) | na28_FW | CACAAAAAAGACCTGATCGCGTACCTGGAGAAAGAAATCGCGAATCTGCGTAAGAAAATCGAAGAATGACAATTAATCATCGGCTCG | pJW61 |
| | na29_FW | AACCTGGTTGCGCAGCTCGAAAACGAAGTTGCGTCTCTGGAAAATGAGAACGAAACCCTGAAGAAAAAGAACCTGCACAAAAAAGACCTGATCGCGTAC | pJW61 |
| | na30_RV | TGAGATAGCTGCAGTCAGCTCG | pJW61 |
| | na06_FW | AACGAGCTGACTGCAGCTATCTCATTGCCTTTATCGTTTGGTCAAC | *X. nematophila* ATCC 19061 |
| | na37_RV | CGTTTTCGAGCTGCGCAACCAGGTTCATGGCTGGCGTTAGTACCG | *X. nematophila* ATCC 19061 |
| pNA18 (NRPS-27) | na32_RV | GTTCTGTTCATCACGTTCCAGCTGCAGGTTGTCTTTTTTCAGACGTGCGATTTTCTTACGCAGATACGCGTTACGCGCCATGGAATTCCTCCTGTTAGCC | pCDF_ara/tacI |
| | na33_RV | CTGTTCGTGAGACGCAACTTCGTTTTCGAGACGCGCGATTTCGTCACGCAGGTTCGCGATGATTTTTTCCAGGTTCTGTTCATCACGTTCCAGC | pCDF_ara/tacI |
| | na34_FW | TGACAATTAATCATCGGCTCG | pCDF_ara/tacI |
| | na38_FW | AAACGAAGTTGCGTCTCACGAACAGTTGCCGCTGATTGATCTCAC | *X. nematophila* ATCC 19061 |
| | na7_RV | CGAGCCGATGATTAATTGTCACAGCGCCTCCACTTCG | *X. nematophila* ATCC 19061 |
| pNA26 (NRPS-34, NRPS-35, NRPS-36, NRPS-37, NRPS-38, NRPS-39, NRPS-40, NRPS-41) | KB-pACYC-II-FW | AACGAGAAGGAGGAATTAAAATCG | pJW61 |
| | KB-pACYC-II-RV | CATGGAATTCCTCCTGTTAGC | pJW61 |
| | na51 | GCTAACAGGAGGAATTCCATGAAAGATAGCATGGCTAAAAAGGAAAT | *P. luminescens* TTO1 |

| | | | |
|---|---|---|---|
| | na52 | CGATTTTAATTCCTCCTTCTCGTTATAAATTTGGCGAGCAAAAGC | *P. luminescens* TTO1 |
| pNA27 (NRPS-31, NRPS-33, NRPS-34, NRPS-35, NRPS-39) | KB-pCOLA-II-FW | TGACAATTAATCATCGGCTCG | pJW62 |
| | KB-pCOLA-II-RV | TGAGATAGCTGCAGTCAGCTCG | pJW62 |
| | na53 | CGAGCTGACTGCAGCTATCTCAGTCGCGCCACAGGGAG | *P. luminescens* TTO1 |
| | na54 | CGTTTTCGAGCTGCGCAACCAGGTTGTAAGCTTGGCGAGCAAAGG | *P. luminescens* TTO1 |
| pNA28 (NRPS-29, NRPS-33, NRPS-34, NRPS-36, NRPS-37) | na33_RV | CTGTTCGTGAGACGCAACTTCGTTTTCGAGACGCGCGATTTCGTCACGCAGGTTCG CGATGATTTTTTCCAGGTTCTGTTCATCACGTTCCAGC | pCDF_ara/tacI |
| | na34_FW | TGACAATTAATCATCGGCTCG | pCDF_ara/tacI |
| | na55 | GAAGTTGCGTCTCACGAACAGCAAGCGCCACAAGGGGA | *P. luminescens* TTO1 |
| | na56 | CGAGCCGATGATTAATTGTCACAGCGCCTCCGCTTCAC | *P. luminescens* TTO1 |
| pNA29 | KB-pACYC-II-FW | AACGAGAAGGAGGAATTAAAATCG | pJW61 |
| | KB-pACYC-II-RV | CATGGAATTCCTCCTGTTAGC | pJW61 |
| | na57 | GCTAACAGGAGGAATTCCATGAAAGGTAGTATTGCTAAAAAGGGAGATG | *X. szentirmaii* DSM16338 |
| | na58 | CGATTTTAATTCCTCCTTCTCGTTATAATGCTGACGGGCAAATG | *X. szentirmaii* DSM16338 |
| pNA30 (NRPS-30, NRPS-36, NRPS-41, NRPS-42) | KB-pCOLA-II-FW | TGACAATTAATCATCGGCTCG | pJW62 |
| | KB-pCOLA-II-RV | TGAGATAGCTGCAGTCAGCTCG | pJW62 |
| | na59 | CGAGCTGACTGCAGCTATCTCAGAATCCCCACAAGGGGAGA | *X. szentirmaii* DSM16338 |
| | na60 | TCGAGCTGCGCAACCAGGTTATAATGCTGACGGGCAAACG | *X. szentirmaii* DSM16338 |
| pNA31 (NRPS-32, NRPS-39, NRPS-40, NRPS-41, NRPS-42) | KB-pACYC-II-FW | AACGAGAAGGAGGAATTAAAATCG | pJW61 |
| | KB-pACYC-II-RV | CATGGAATTCCTCCTGTTAGC | pJW61 |
| | na61 | AAAACGAAGTTGCGTCTCACGAACAGGAGTTGCCACAAGGTGAAA | *X. szentirmaii* DSM16338 |
| | na62 | CGAGCCGATGATTAATTGTCAAATATTATTTACTATATTGTATTCCTCTGTACCA | *X. szentirmaii* DSM16338 |
| pNA35 | KB-pCOLA-II-FW | TGACAATTAATCATCGGCTCG | pJW62 |
| | KB-pCOLA-II-RV | TGAGATAGCTGCAGTCAGCTCG | pJW62 |
| | na68 | TAACGAGCTGACTGCAGCTATCTCAGAAACTCCGGTCGGTAAAGTAG | *X. bovienii* SS2004 |
| | na69 | TTCGTTTTCGAGCTGCGCAACCAGGTTATAGCCGCGCACCACTAC | *X. bovienii* SS2004 |
| pNA59 (NRPS-47, NRPS-48) | KB-pACYC-FW | GAACAGTTAAAACAGAAGCGTGAACAATTAAAGCAAAAGATCGCCAATCTGCGTAA GGAGATCGAAGCCTACAAGTGACAATTAATCATCGGCTCG | pCK_0402 |
| | KB-pACYC-RV | TTCACGCTTCTGTTTTAACTGTTCGATGCGATTACGCAATTCAGCCTTTTTCGATTTT AATTCCTCCTTCTCGTTCATGGAATTCCTCCTGTTAGC | pCK_0402 |
| | na125_FW | GCTAACAGGAGGAATTCCATGGATAACATTCTGGCCTCG | *X. bovienii* SS2004 |
| | na126_RV | CAGATTTTAAACAGAGCCGCTATGTTTATAACGAGAAGGAGGAATTAAAATCG | *X. bovienii* SS2004 |
| pNA60 (NRPS-47) | KB-pCOLA-FW | CATTGACAAAGAGCTGCGTGCCAACGAAAACGAACTTCGCGCCCTTGATAACGAGC TGACTGCAGCTATCTCATGACAATTAATCATCGGCTCG | pCOLA_ara/tacI |
| | KB-pCOLA-RV | TTGGCACGCAGCTCTTTGTCAATGGCATTTAACTCGCGGTCCAAGGCTTTCAGTTCA CGCTCTTCAGCATAGAACATGGAATTCCTCCTGTTAGC | pCOLA_ara/tacI |
| | na127_FW | AACGAGCTGACTGCAGCTATCTCACGTGCCGCAGAAACGG | *X. bovienii* SS2004 |
| | na128_RV | ATACGAGCCGATGATTAATTGTCAATCCACCAGCTCCAACAC | *X. bovienii* SS2004 |
| pCOLA_ara_xtpS _tacI_JW | jw0136_FW | CGCTGCTGGTTCTGGCGATTGACAATTAATCATCGGCTCG | pCOLA_ara/tacI |
| | jw0137_RV | AACGGGTATGGAGAAACAGTAGAGAGTTGCGATAAAAAGCG | pCOLA_ara/tacI |
| | AL-GxpS-2-1 | ACTGTTTCTCCATACCCGTTTTTTTGGGCTAACAGGAGGAATTCCATGAAAGATAGC ATGGCTAAAAAGG | *X. nematophila* ATCC 19061 |
| | AD64 | TCGCCAGAACCAGCAGCGGAGCCAGCGGATCCGGCGCGCCTTACAGCGCCTCCA C | *X. nematophila* ATCC 19061 |
| pCOLA_ara_gxpS _tacI_JW | JW_tacI_PstI_FW 2 | CTGCAGGAGCTGTTGACAAT | pCOLA_ara/tacI |
| | jw0064_RV | CATGGAATTCCTCCTGTTAGCC | pCOLA_ara/tacI |
| | jw0124_FW | GGGCTAACAGGAGGAATTCCATGAAAGATAGCATGGCTAAAAAGGAAATTATC | *P. luminescens* TTO1 |
| | jw0160_RW | GATTAATTGTCAACAGCTCCTGCAGGCAGATAGAGACGTTTGTTGGC | *P. luminescens* TTO1 |
| | jw0151_FW/ | GCCAACAAACGTCTCTATCTGCTGGATGAACACCG | *P. luminescens* TTO1 |
| | jw0161_RV | GATTAATTGTCAACAGCTCCTGCAGTCACAGCGCCTCCGCTTCAC | *P. luminescens* TTO1 |

### 3 Supplementary Figures

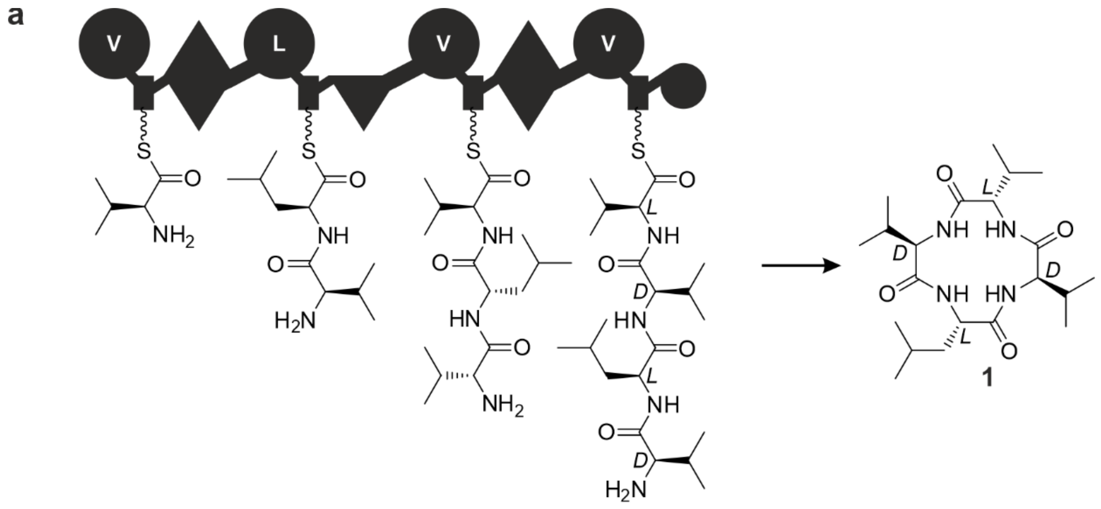

**Figure S1.** A schematic representation of the xenotetrapeptide (**1**) producing NRPS (XtpS).

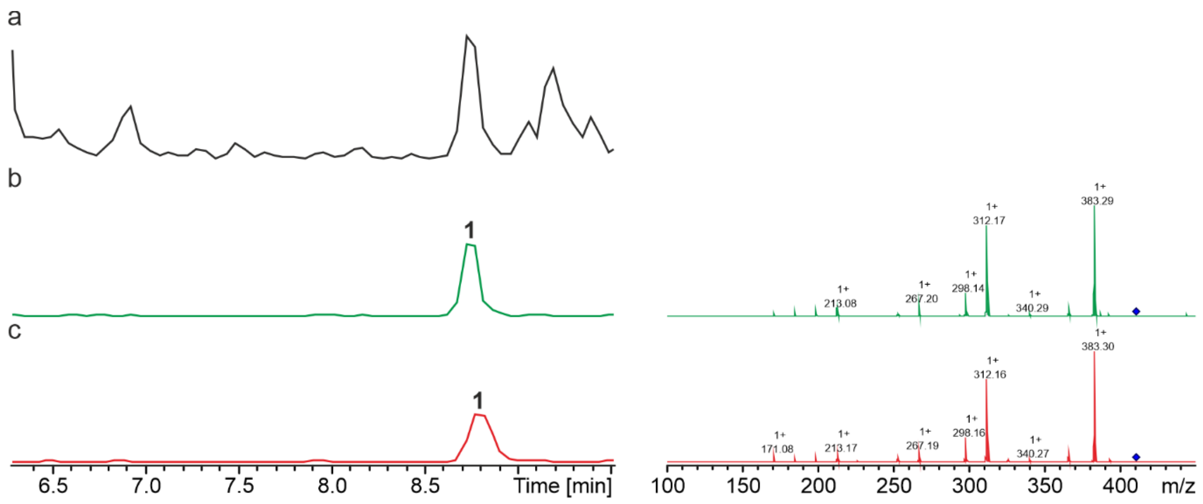

**Figure S2.** HPLC/MS data refers to Figure 2a (NRPS-1-4) of compound **1** produced in *E. coli* DH10B::*mtaA*. (a) Base Peak Chromatogram (BPC) of an exemplary culture extract. (b) EIC/MS$^2$ of **1** (*m/z* $[M+H]^+$ = 411.30). (c) EIC/MS$^2$ data of synthetic **1** (*m/z* $[M+H]^+$ = 411.30).

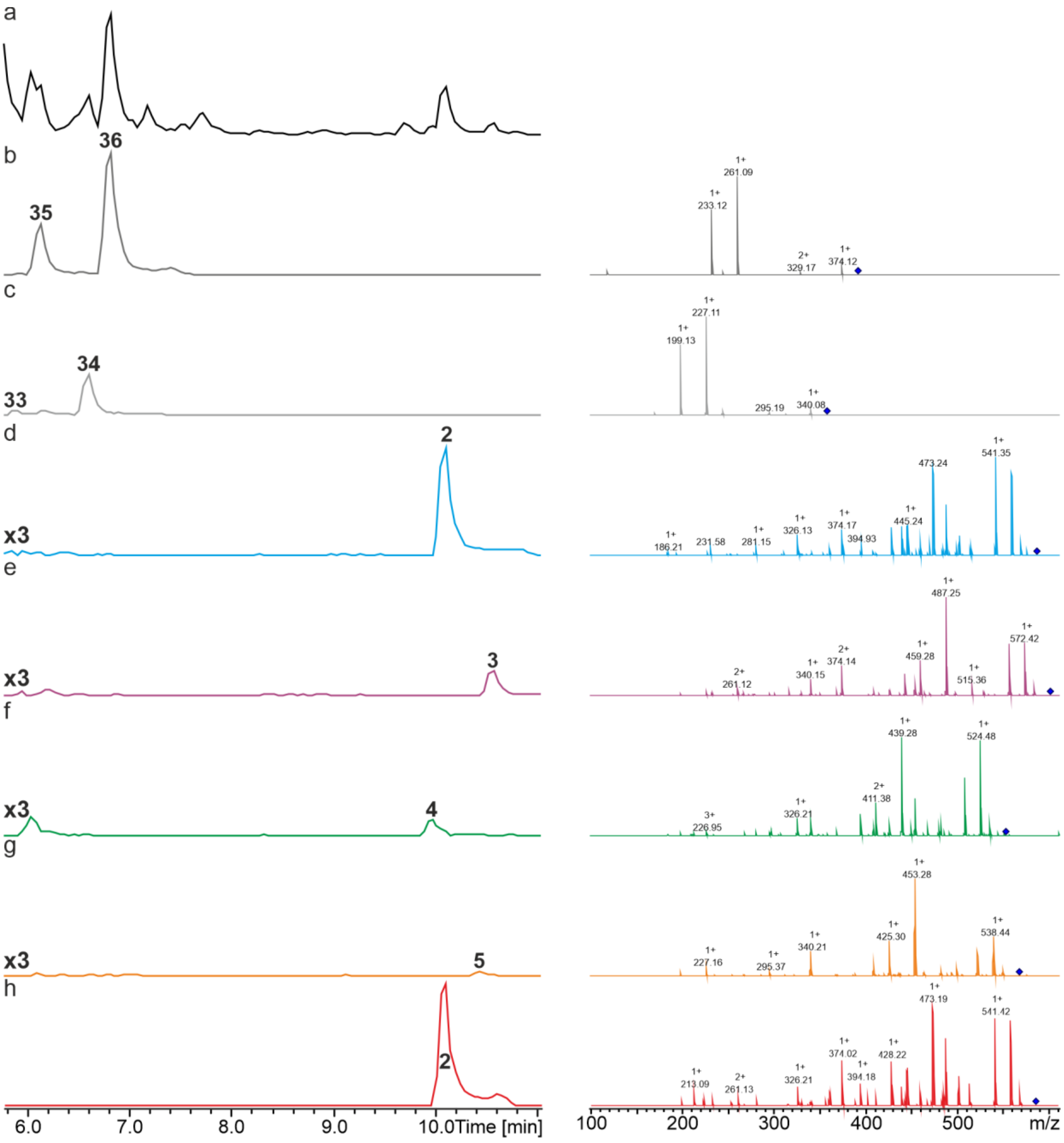

**Figure S3.** HPLC/MS data refers to Figure 2b (NRPS-5) of compounds **2-5**, **33**/**34** and **35**/**36** produced in *E. coli* DH10B::*mtaA*. (a) Base Peak Chromatogram (BPC) of an exemplary culture extract. (b) Extracted ion chromatogram (EIC)/MS$^2$ of **35/36** (*m/z* [M+H]$^+$ = 392.25). (c) EIC/MS$^2$ of **33/34** (*m/z* [M+H]$^+$ = 358.27). (d) EIC/MS$^2$ of **2** (*m/z* [M+H]$^+$ = 586.40). (e) EIC/MS$^2$ of **3** (*m/z* [M+H]$^+$ = 600.41). (f) EIC/MS$^2$ of **4** (*m/z* [M+H]$^+$ = 552.41). (g) EIC/MS$^2$ **5** (*m/z* [M+H]$^+$ = 566.43). EICs of **2-5** are depicted with threefold increased intensity. (h) EIC/MS$^2$ of synthetic **2** (*m/z* [M+H]$^+$ = 586.40).

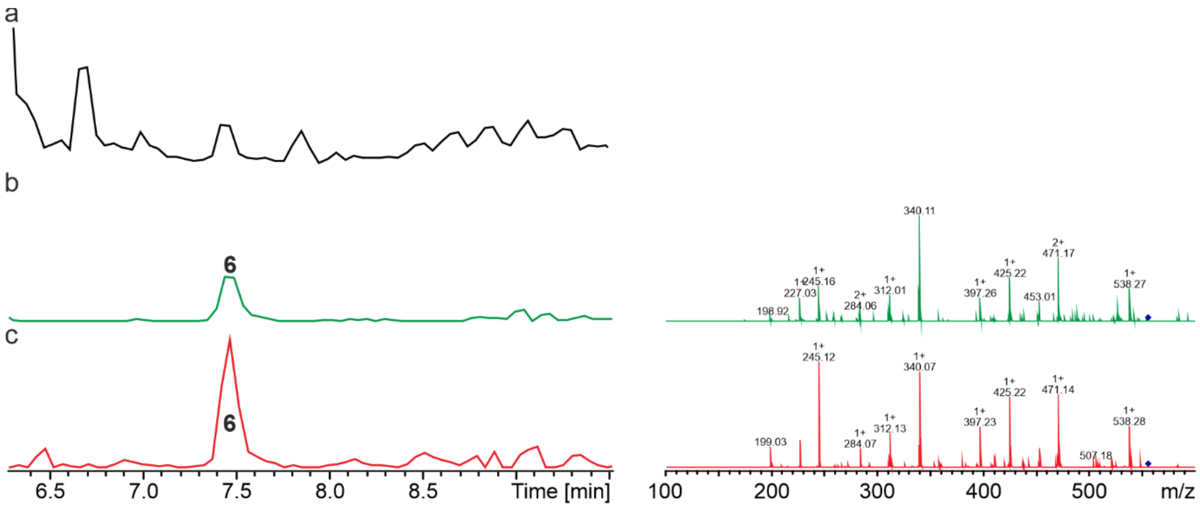

**Figure S4.** HPLC/MS data refers to Figure 2b (NRPS-6) of compound **6** produced in *E. coli* DH10B::*mtaA*. (a) Base Peak Chromatogram (BPC) of an exemplary culture extract. (b) Extracted ion chromatogram (EIC)/MS$^2$ of **6** (*m/z* $[M+H]^+$ = 556.35). (c) EIC/MS$^2$ of synthetic **6** (*m/z* $[M+H]^+$ = 556.35).

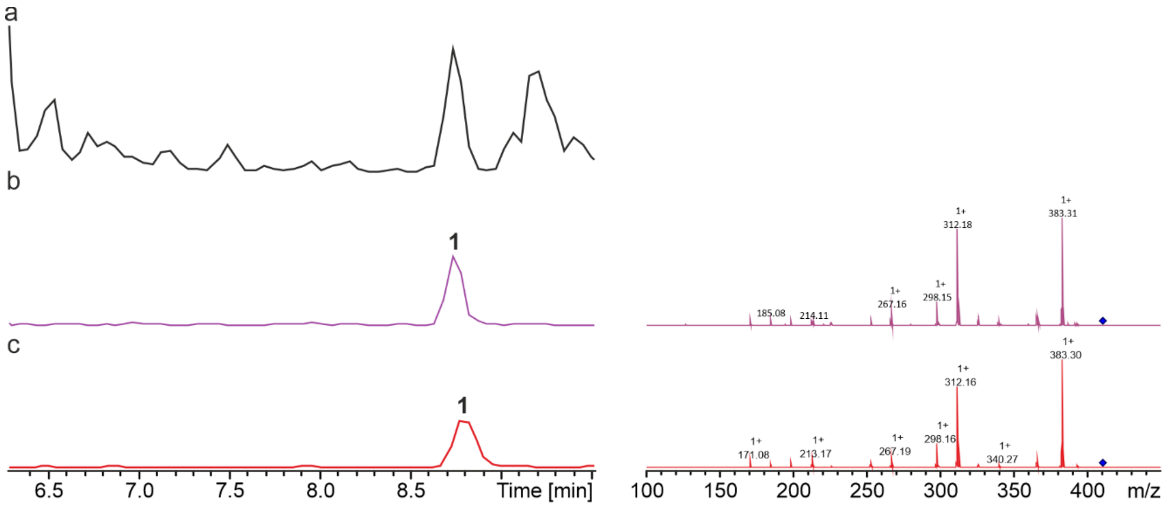

**Figure S5.** HPLC/MS data refers to Figure 3a (NRPS-7) of compound **1** produced in *E. coli* DH10B::*mtaA*. (a) Base Peak Chromatogram (BPC) of an exemplary culture extract. (b) Extracted ion chromatogram (EIC)/MS$^2$ of **1** (*m/z* $[M+H]^+$ = 411.30). (c) EIC/MS$^2$ of synthetic **1** (*m/z* $[M+H]^+$ = 411.30).

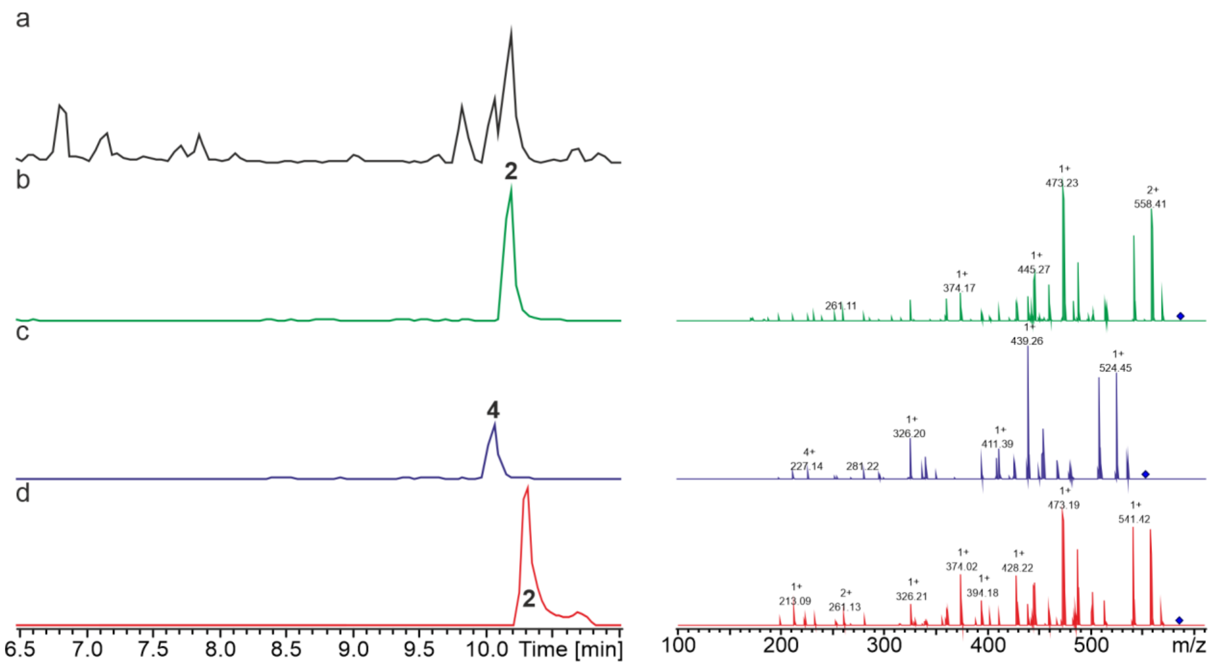

**Figure S6.** HPLC/MS data refers to Figure 3a (NRPS-8) of compounds **2** and **4** produced in *E. coli* DH10B::*mtaA*. (a) Base Peak Chromatogram (BPC) of an exemplary culture extract. (b) Extracted ion chromatogram (EIC)/MS$^2$ of **2** ($m/z$ [M+H]$^+$ = 586.40). (c) Extracted ion chromatogram (EIC)/MS$^2$ of **4** ($m/z$ [M+H]$^+$ = 552.41). (d) EIC/MS$^2$ of synthetic **2** ($m/z$ [M+H]$^+$ = 586.40).

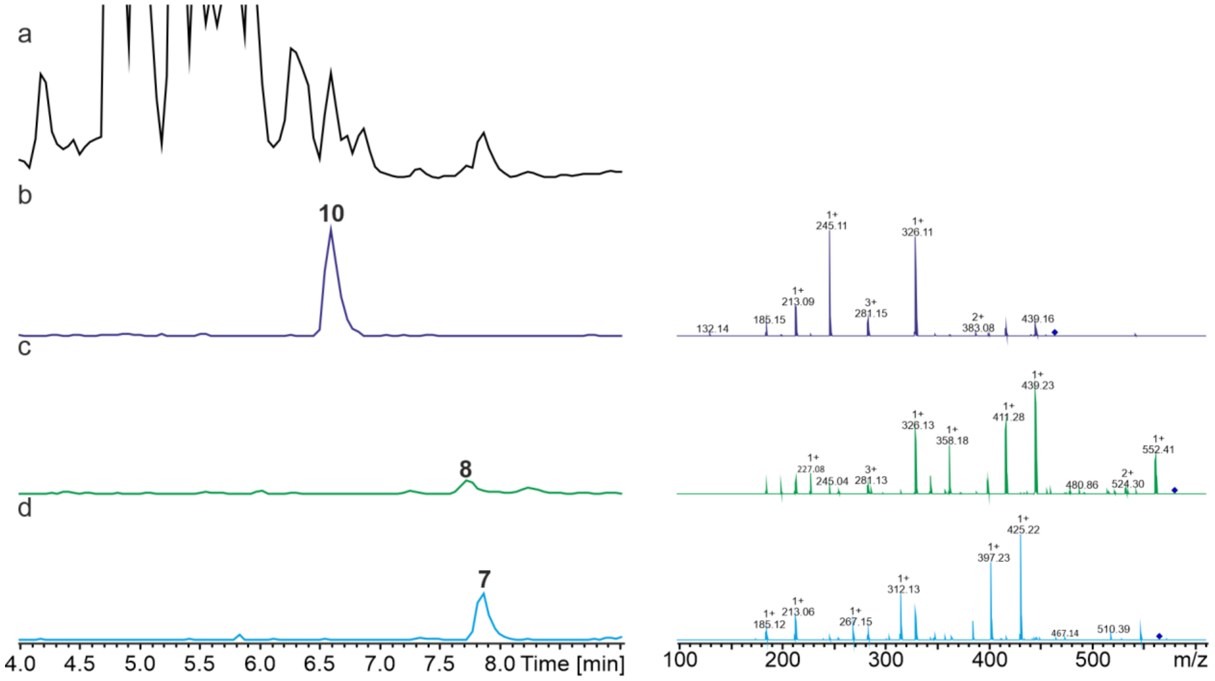

**Figure S7.** HPLC/MS data refers to Figure 3b (NRPS-9) of compounds **7**, **8** and **10** produced in *E. coli* DH10B::*mtaA.* (a) Base Peak Chromatogram (BPC) of an exemplary culture extract. (b) Extracted ion chromatogram (EIC)/MS$^2$ of **10** ($m/z$ [M+H]$^+$ = 457.34). (c) Extracted ion chromatogram (EIC)/MS$^2$ of **8** ($m/z$ [M+H]$^+$ = 570.42). (d) Extracted ion chromatogram (EIC)/MS$^2$ of **7** ($m/z$ [M+H]$^+$ = 556.41).

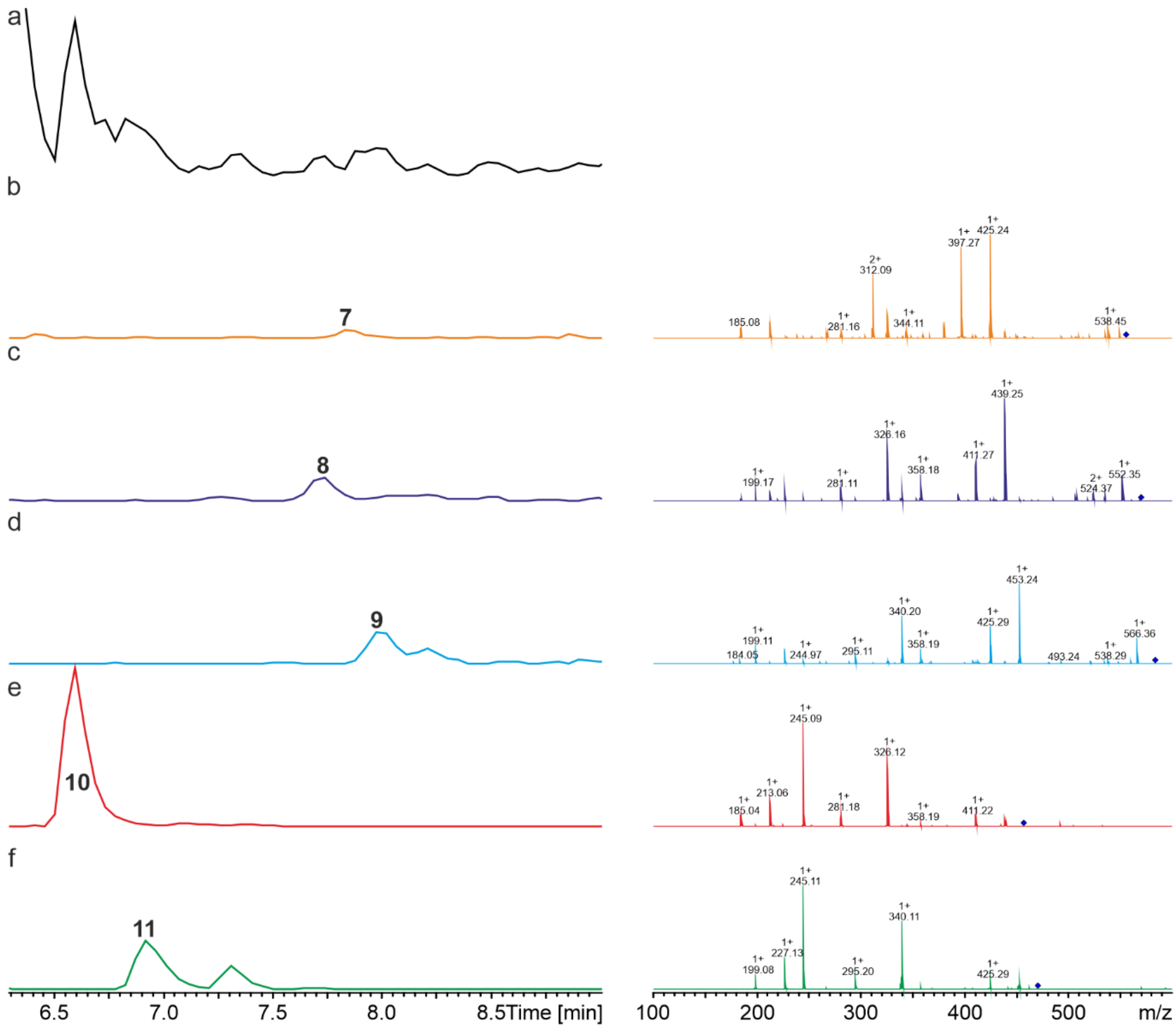

**Figure S8.** HPLC/MS data refers to Figure 3b (NRPS-10) of compounds **7-11** produced in *E. coli* DH10B::*mtaA*. (a) Base Peak Chromatogram (BPC) of an exemplary culture extract. (b) Extracted ion chromatogram (EIC)/MS$^2$ of **7** ($m/z$ [M+H]$^+$ = 556.41). (c) Extracted ion chromatogram (EIC)/MS$^2$ of **8** ($m/z$ [M+H]$^+$ = 570.42). (d) Extracted ion chromatogram (EIC)/MS$^2$ of **9** ($m/z$ [M+H]$^+$ = 584.44). (e) Extracted ion chromatogram (EIC)/MS$^2$ of **10** ($m/z$ [M+H]$^+$ = 457.34). (f) Extracted ion chromatogram (EIC)/MS$^2$ of **11** ($m/z$ [M+H]$^+$ = 471.35).

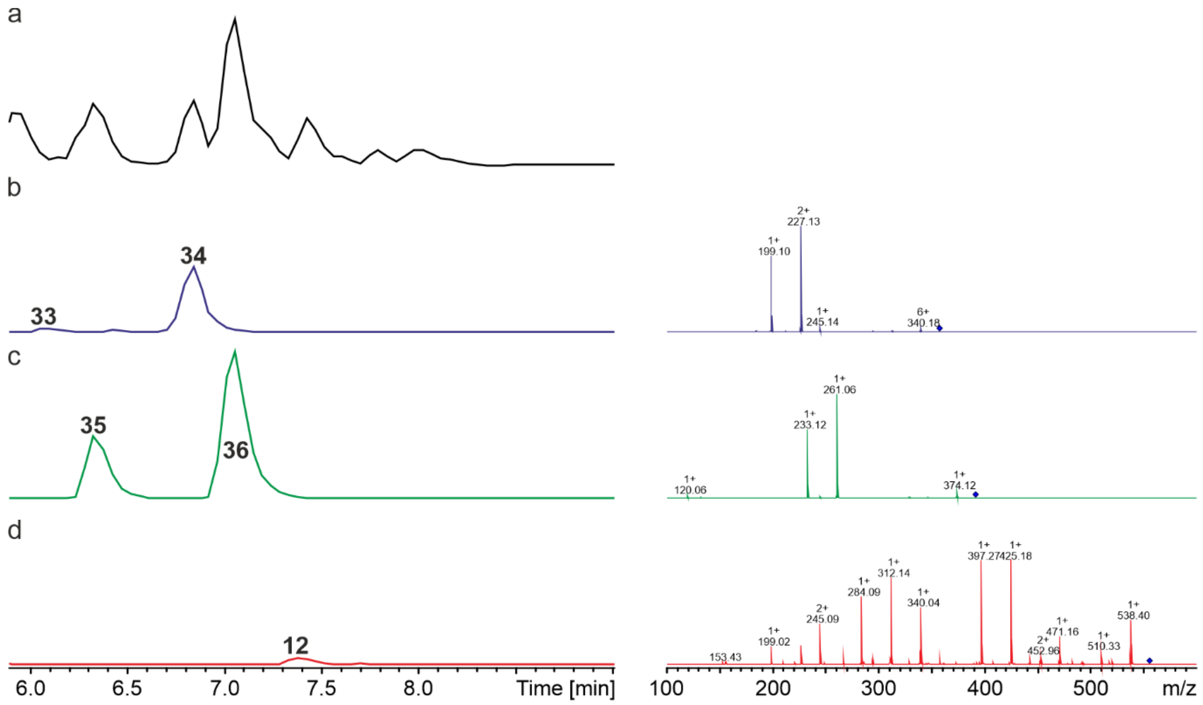

**Figure S9.** HPLC/MS data refers to Figure 3b (NRPS-11) of compounds **33**/**34**, **35**/**36** and **12** produced in *E. coli* DH10B::*mtaA*. (a) Base Peak Chromatogram (BPC) of an exemplary culture extract. (b) EIC/MS$^2$ data data of **33**/**34** (*m/z* $[M+H]^+$ = 358.27). (c) EIC/MS$^2$ data of **35/36** (*m/z* $[M+H]^+$ = 392.25). (d) Extracted ion chromatogram (EIC)/MS$^2$ of **12** (*m/z* $[M+H]^+$ = 556.35).

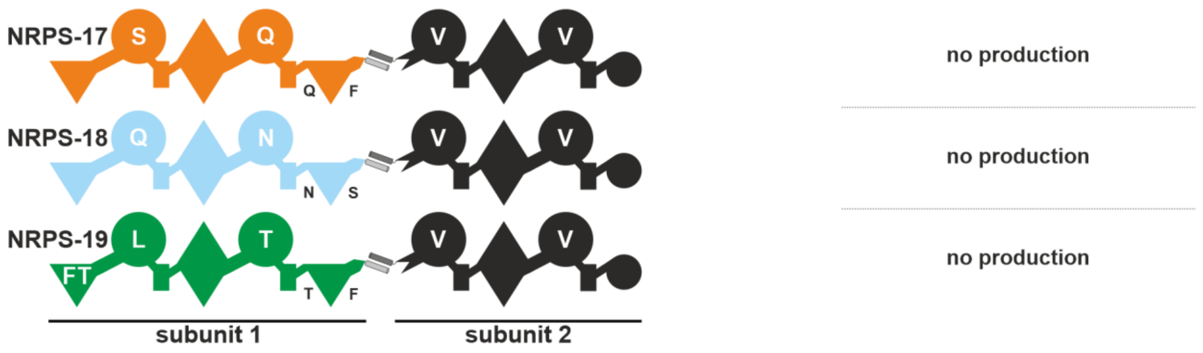

**Figure S10.** A schematic representation of non-functional recombinant type S NRPSs using subunit 1 building blocks from AmbS XldS and SzeS combined with XtpS subunit 2.

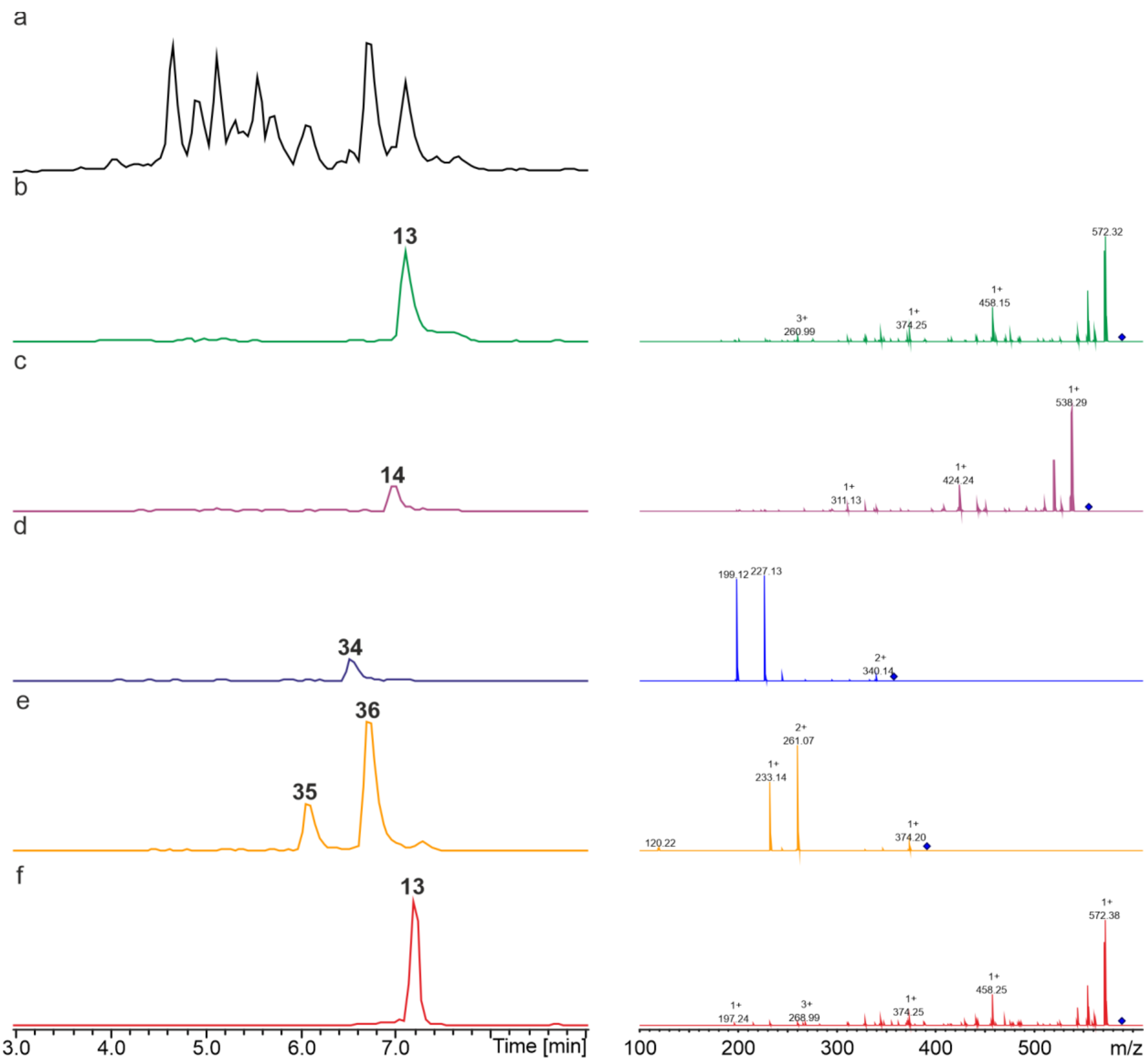

**Figure S11.** HPLC/MS data refers to Figure 3c (NRPS-13) of compounds **13**, **14**, **33**/**34** and **35**/**36** produced in *E. coli* DH10B::*mtaA*. (a) Base Peak Chromatogramm (BPC) of an exemplary culture extract. (b) EIC/MS$^2$ data of **13** (*m/z* [M+H]$^+$ = 589.33). (c) EIC/MS$^2$ data of **14** (*m/z* [M+H]$^+$ = 555.35). (d) EIC/MS$^2$ data data of **34** (*m/z* [M+H]$^+$ = 358.27). (e) EIC/MS$^2$ data of **35**/**36** (*m/z* [M+H]$^+$ = 392.25). (f) EIC/MS$^2$ data of synthetic **13** (*m/z* [M+H]$^+$ = 589.33).

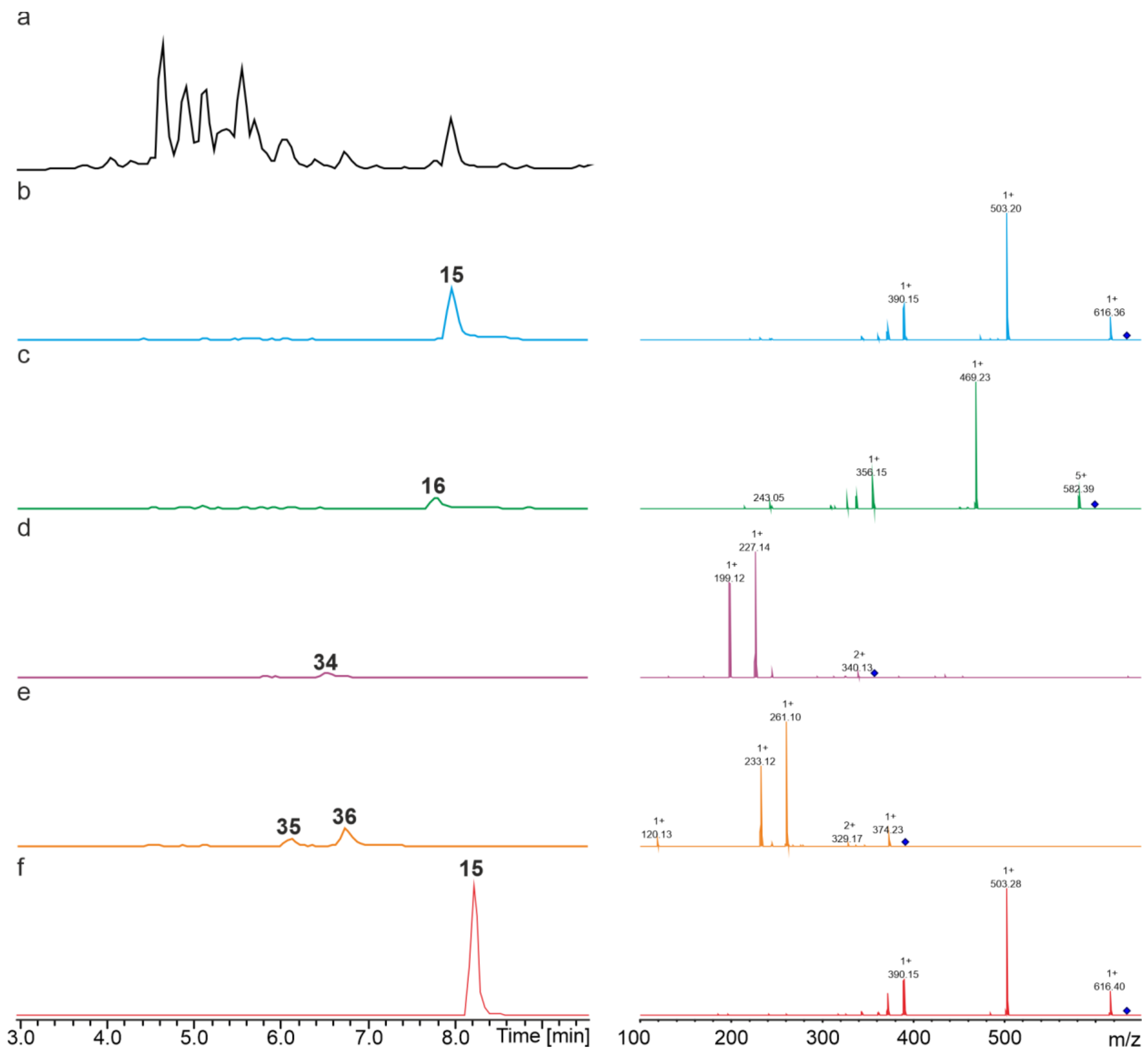

**Figure S12.** HPLC/MS data refers to Figure 3c (NRPS-14) of compounds **15**, **16**, **34** and **35**/**36** produced in *E. coli* DH10B::*mtaA*. (a) Base Peak Chromatogramm (BPC) of an exemplary culture extract. (b) EIC/MS$^2$ data of **15** (*m/z* $[M+H]^+$ = 634.38). (c) EIC/MS$^2$ data of **16** (*m/z* $[M+H]^+$ = 600.40). (d) EIC/MS$^2$ data of **34** (*m/z* $[M+H]^+$ = 358.27). (e) EIC/MS$^2$ data of **35/36** (*m/z* $[M+H]^+$ = 392.25). (f) EIC/MS$^2$ data of synthetic **15** (*m/z* $[M+H]^+$ = 634.38).

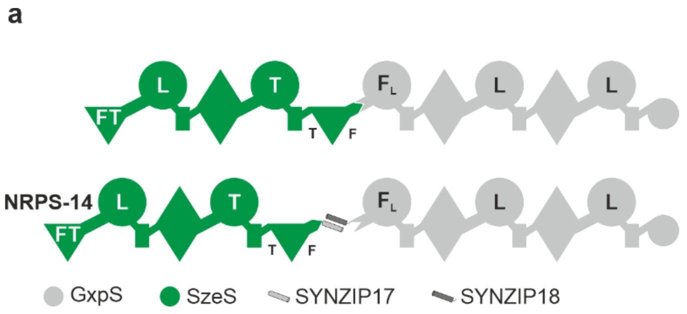

| Peptide | Production (mg $l^{-1}$) |
|---|---|
| **15** | 14.73±0.45 |
| **16** | 1.68±0.71 |
| **15** | 23.82±2.64 |
| **16** | 4.07±1.10 |

**Figure S13.** Comparison of production yields of a homologous *in cis* and *trans* NRPS-14. The colour code of the NRPS subunits is depicted at the bottom of the figure. The domain assignment is as in Figure 1 plus FT (formyltransferase, N-terminal triangle) and specificities are assigned for the entire A domains.

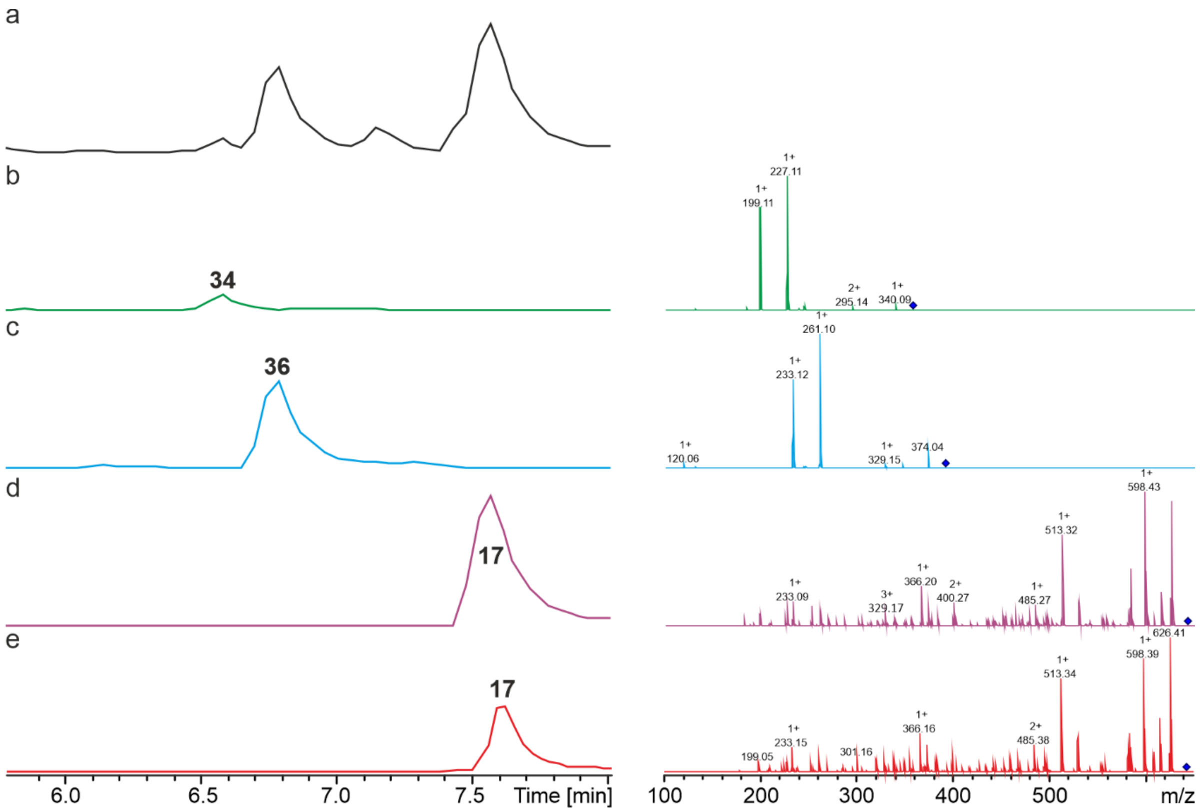

**Figure S14.** HPLC/MS data refers to Figure 3d (NRPS-15) of compounds **34**, **36** and **17** produced in *E. coli* DH10B::*mtaA*. (a) Base Peak Chromatogramm (BPC) of an exemplary culture extract. (b) EIC/MS$^2$ data of **34** (*m/z* $[M+H]^+$ = 358.27). (c) EIC/MS$^2$ data of **36** (*m/z* $[M+H]^+$ = 392.25). (d) EIC/MS$^2$ data of **17** (*m/z* $[M+H]^+$ = 643.43). (e) EIC/MS$^2$ data of synthetic **17** (*m/z* $[M+H]^+$ = 643.43).

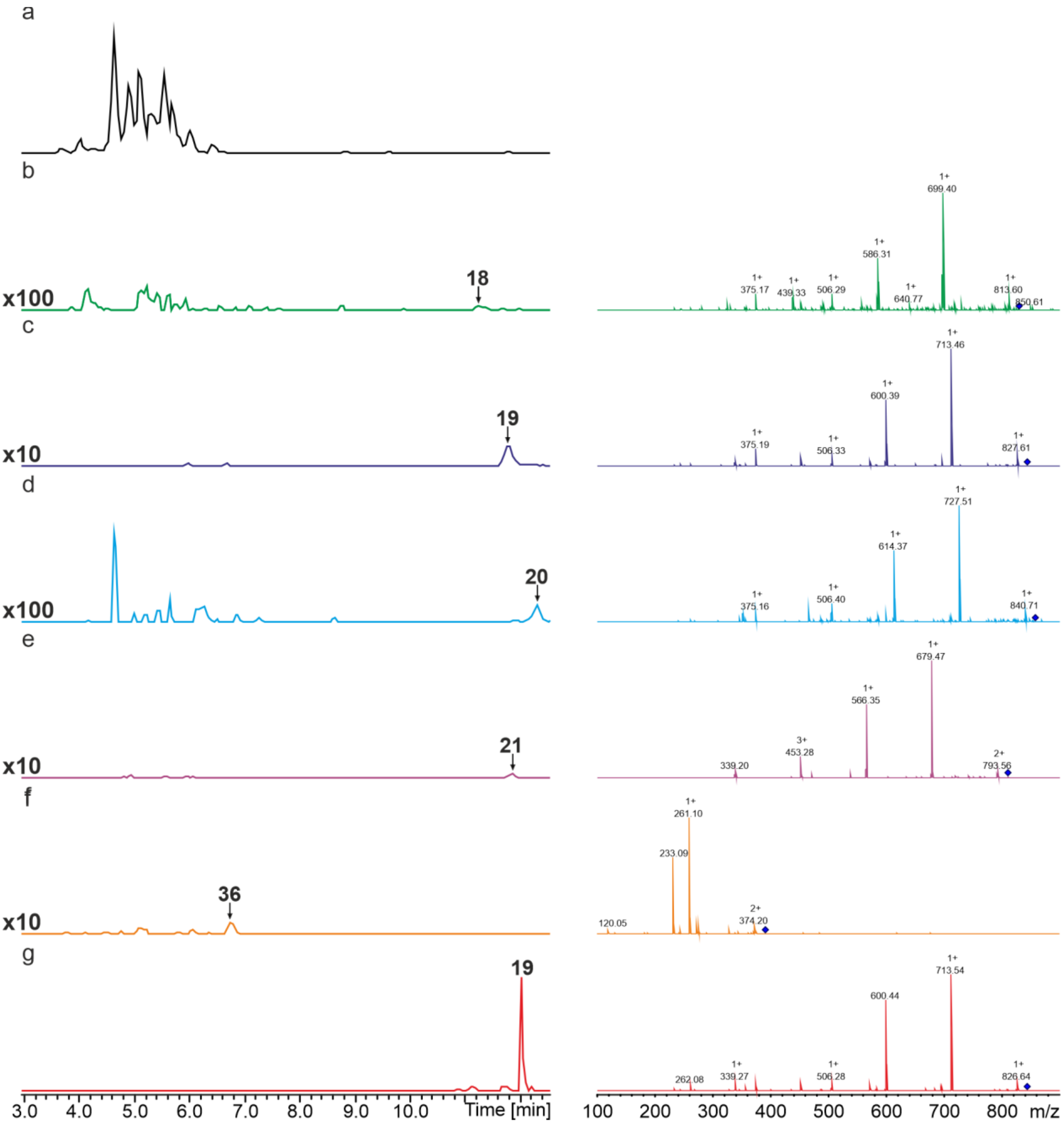

**Figure S15.** HPLC/MS data refers to Figure 3d (NRPS-16) of compounds **18-21** and **36** produced in *E. coli* DH10B::*mtaA*. (a) Base Peak Chromatogramm (BPC) of an exemplary culture extract. (b) EIC/MS$^2$ data of **18** (*m/z* [M+H]$^+$ = 830.54). (c) EIC/MS$^2$ data of **19** (*m/z* [M+H]$^+$ = 844.55). (d) EIC/MS$^2$ data of **20** (*m/z* [M+H]$^+$ = 858.57). (e) EIC/MS$^2$ data of **21** (*m/z* [M+H]$^+$ = 810.57). (f) EIC/MS$^2$ data of **36** (*m/z* [M+H]$^+$ = 392.25). (g) EIC/MS$^2$ data of synthetic **19** (*m/z* [M+H]$^+$ = 844.55).

**a**

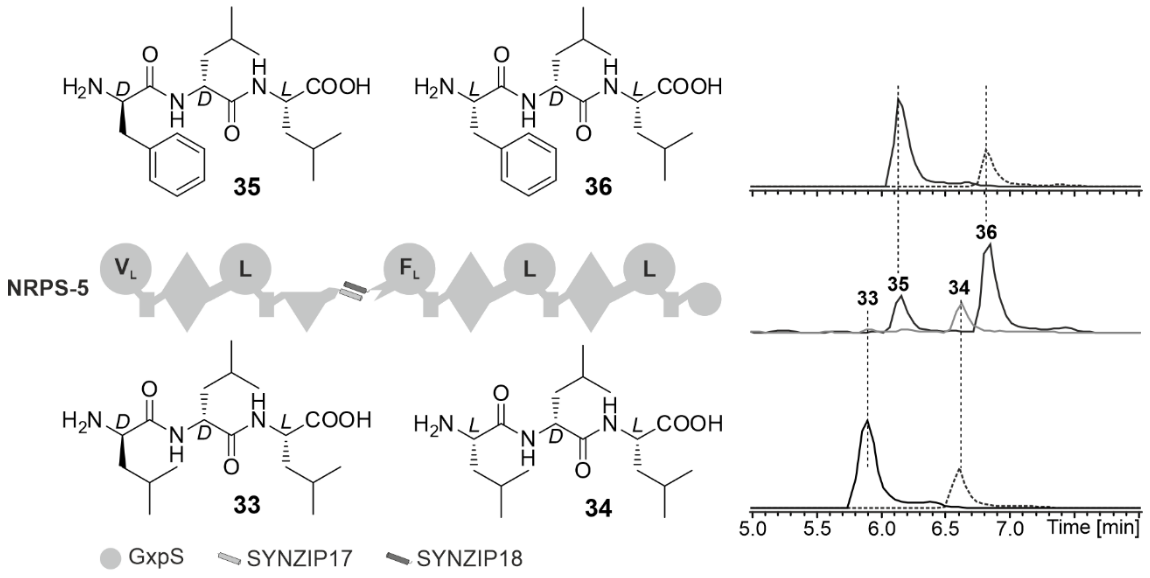

**b**

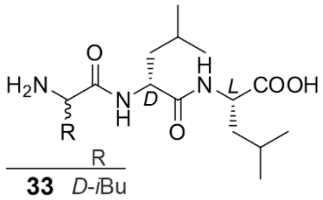

| | R |
|---|---|
| **33** | *D*-*i*Bu |
| **34** | *L*-*i*Bu |
| **35** | *D*-Ph |
| **36** | *L*-Ph |

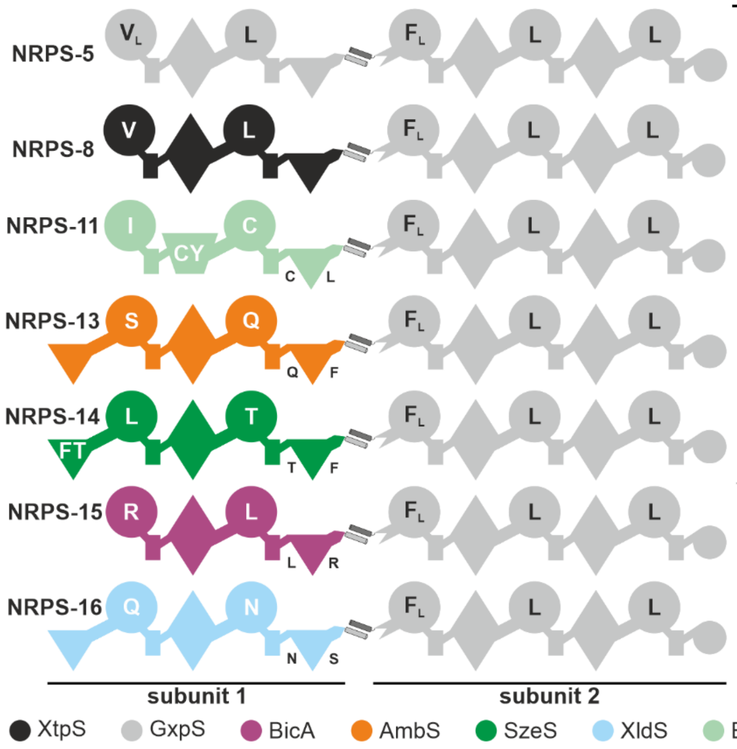

| Peptide | Production (mg $l^{-1}$) |
|---|---|
| 33 | - |
| 34 | 3.59±0.11 |
| 35 | 5.31±0.06 |
| 36 | 16.08±0.19 |
| 33 | 0.06±0.01 |
| 34 | 0.72±0.05 |
| 35 | 1.44±0.09 |
| 36 | 4.92±0.46 |
| 33 | 1.08±0.28 |
| 34 | 20.00±1.18 |
| 35 | 18.25±0.53 |
| 36 | 47.52±2.08 |
| 33 | 0.23±0.06 |
| 34 | 5.11±0.55 |
| 35 | 12.94±0.73 |
| 36 | 37.66±2.36 |
| 33 | 0.10±0.00 |
| 34 | 1.39±0.17 |
| 35 | 1.95±0.28 |
| 36 | 5.62±1.22 |
| 33 | 0.20±0.06 |
| 34 | 10.24±0.56 |
| 35 | 2.31±0.04 |
| 36 | 86.23±2.80 |
| 33 | - |
| 34 | - |
| 35 | 0.15±0.02 |
| 36 | 0.44±0.03 |

**Figure S16.** (a) Production of D/L-tripeptides exemplary of NRPS-5. The tripeptide production is related to the unpaired activity of GxpS subunit 2 resulted in the production of peptides **33/34** and **35/36**. The different epimers could be identified by their retention times. (b) Tripeptide **33/34** and **35/36** amounts and yields (determined in triplicates (n=3)) are given. The colour code of the NRPS subunits is depicted at the bottom of the figures. The domain assignment is as in Figure 1.

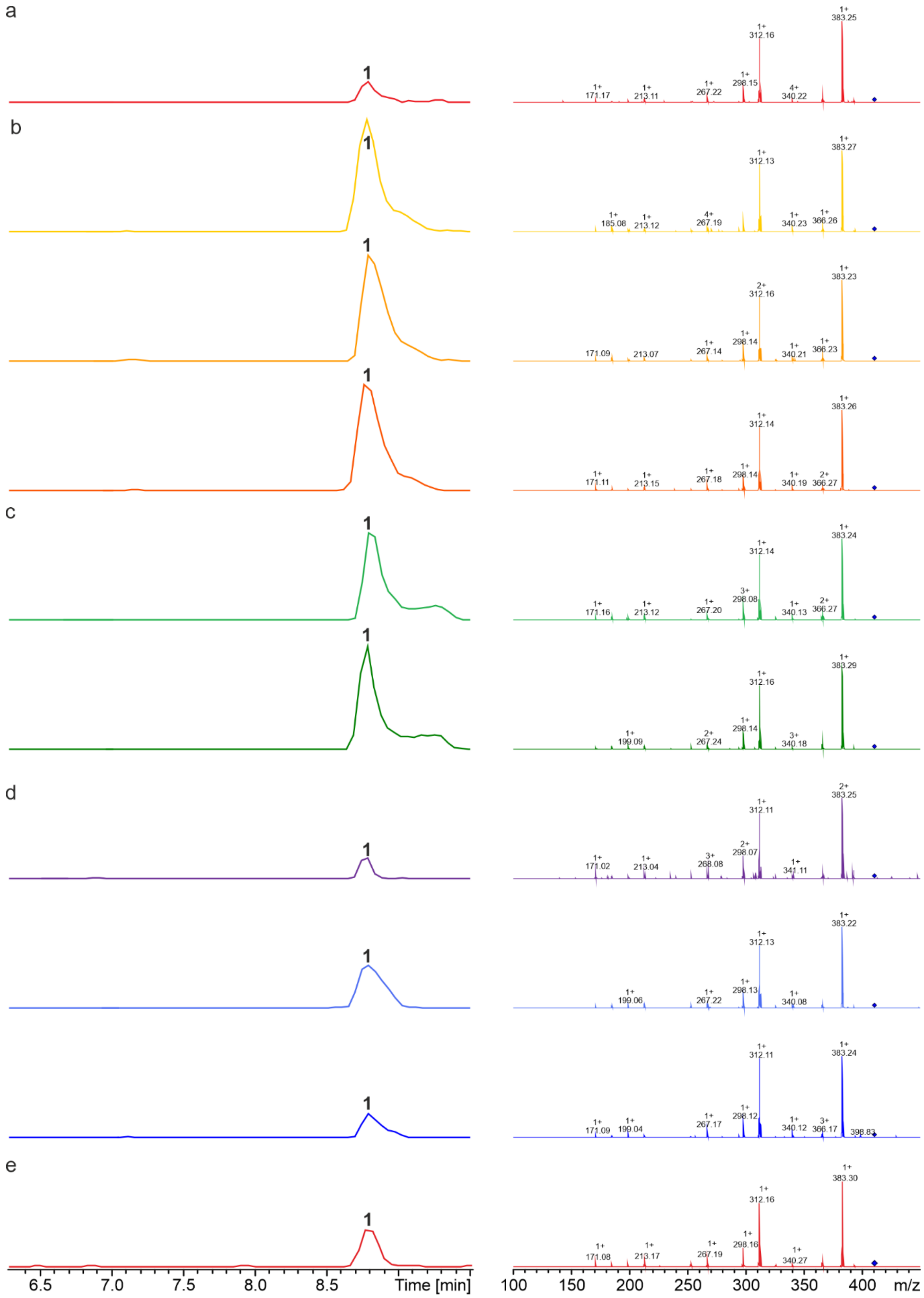

**Figure S17.** HPLC/MS data refers to Figure 4a (NRPS-20), Figure 4b (NRPS-21-23), Figure 4c (NRPS-24 and NRPS-25) and Figure 5 (NRPS-26-28) of compound **1** produced in *E. coli* DH10B::*mtaA*. (a) EIC/MS$^2$ (NRPS-20) of **1** (*m/z* $[M+H]^+$ = 411.30). (b) EIC/MS$^2$ (NRPS-21-23) of **1** (*m/z* $[M+H]^+$ = 411.30). (c) EIC/MS$^2$ (NRPS-24 and NRPS-25) of **1** (*m/z* $[M+H]^+$ = 411.30). (d) EIC/MS$^2$ (NRPS-26-28) of **1** (*m/z* $[M+H]^+$ = 411.30). (e) EIC/MS$^2$ data of synthetic **1** (*m/z* $[M+H]^+$ = 411.30).

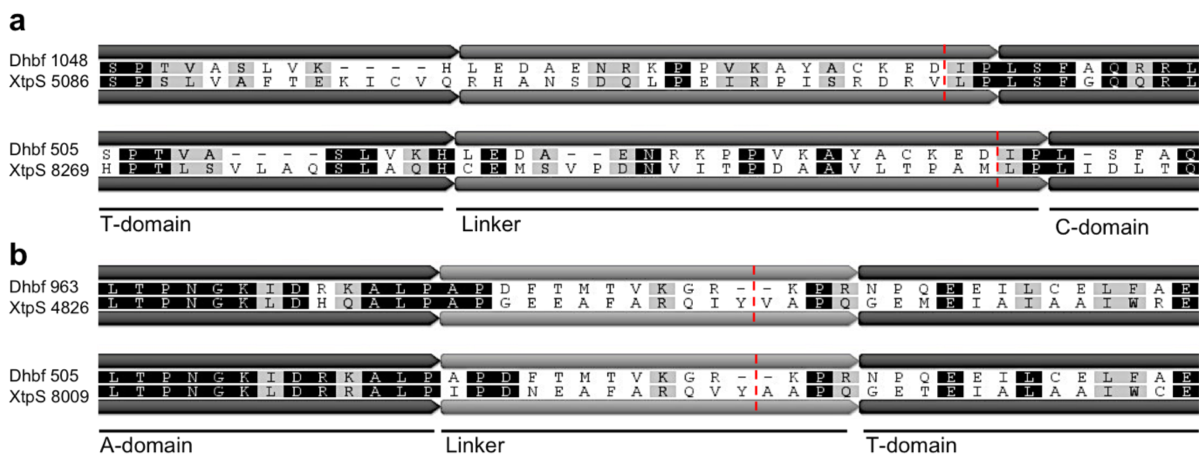

**Figure S18.** Sequence alignments of XtpS linker sequences. (a) XtpS $T_2$-$C_2$ (XtpS 5086) and $T_3$-$C_3$ (XtpS 8269) linker were aligned to the linker region excised from Dhbf crystal structure (Protein Database ID: 5u89). (b) XtpS $A_2$-$T_2$ (XtpS 4826) and $A_3$-$T_3$ (XtpS 8009) linker were aligned to the linker region excised from Dhbf crystal structure. Red dashed lines indicate SYNZIP insertion points.

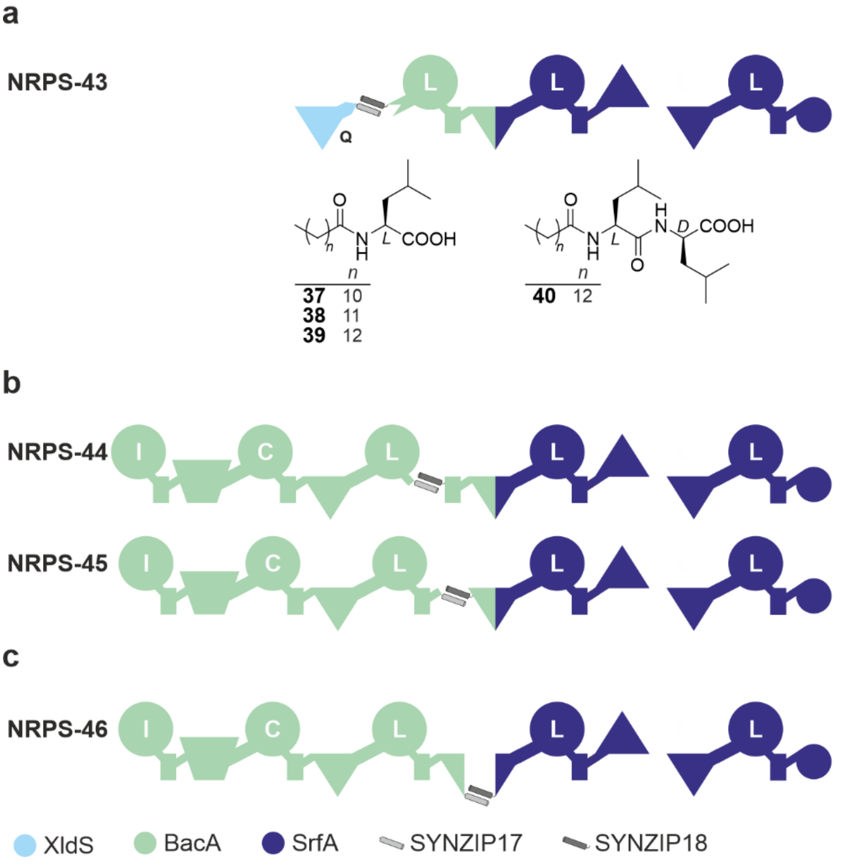

a

| Peptide | Production (mg l$^{-1}$) | % of WT level |
|---|---|---|
| (12:0)-L 37 | - | - |
| (13:0)-L 38 | - | - |
| (14:0)-L 39 | - | - |
| (14:0)-LI 40 | - | - |

b

| Peptide | Production (mg l$^{-1}$) | % of WT level |
|---|---|---|
| IC*LIL | 20.42±2.02 | 127.9 |

NRPS-45: no production

c

| Peptide | Production (mg l$^{-1}$) | % of WT level |
|---|---|---|

NRPS-46: no production

**Figure S19.** Further examples of two component type S NRPS split in between and within RtpS modules.

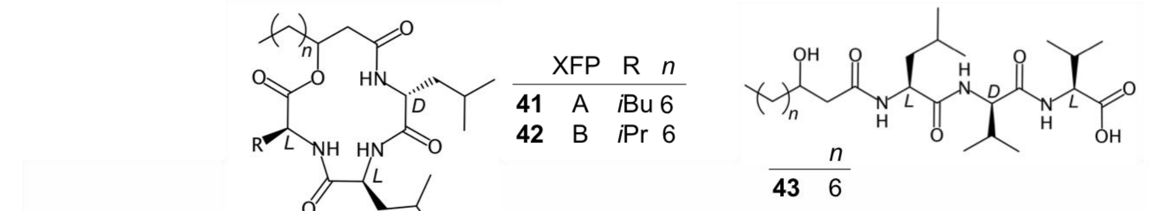

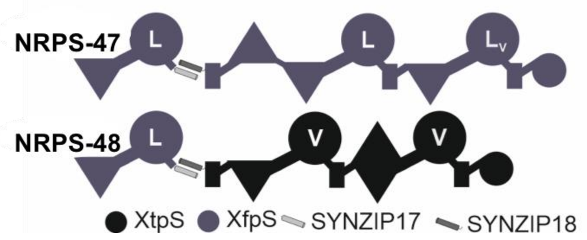

| Peptide | Production (mg l$^{-1}$) | % of WT level |
|---|---|---|
| 10:0(β-hydroxy)-LLL 41 | 0.32±0.03 | 26.69±2.70 |
| 10:0(β-hydroxy)-LLV 42 | 0.35±0.08 | 73.00±16.78 |
| 10:0(β-hydroxy)-LVV 43 | 0.06±0.02 | 13.70±5.07 |

**Figure S20.** Further examples of two component type S NRPS split within modules.

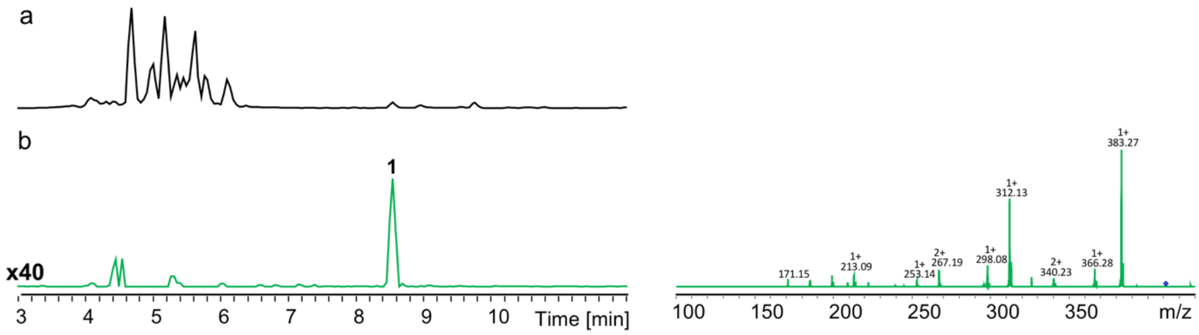

**Figure S21.** HPLC/MS data refers to Figure 6 (NRPS-28) of compound **1** produced in *E. coli* DH10B::*mtaA*. (a) Base Peak Chromatogram (BPC) of an exemplary culture extract. (b) EIC/MS$^2$ data of **1** (*m/z* $[M+H]^+$ = 411.30).

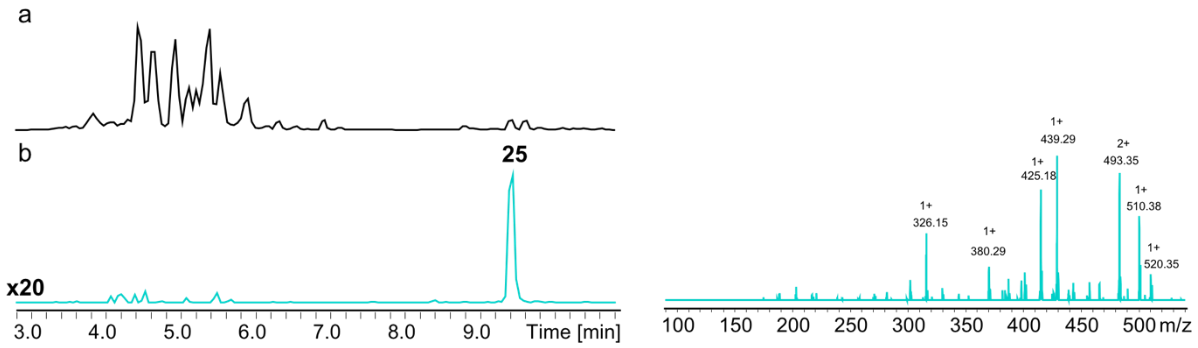

**Figure S22.** HPLC/MS data refers to Figure 6 (NRPS-29) of compound **25** produced in *E. coli* DH10B::*mtaA*. (a) Base Peak Chromatogram (BPC) of an exemplary culture extract. (b) EIC/MS$^2$ data of **25** (*m/z* $[M+H]^+$ = 538.40).

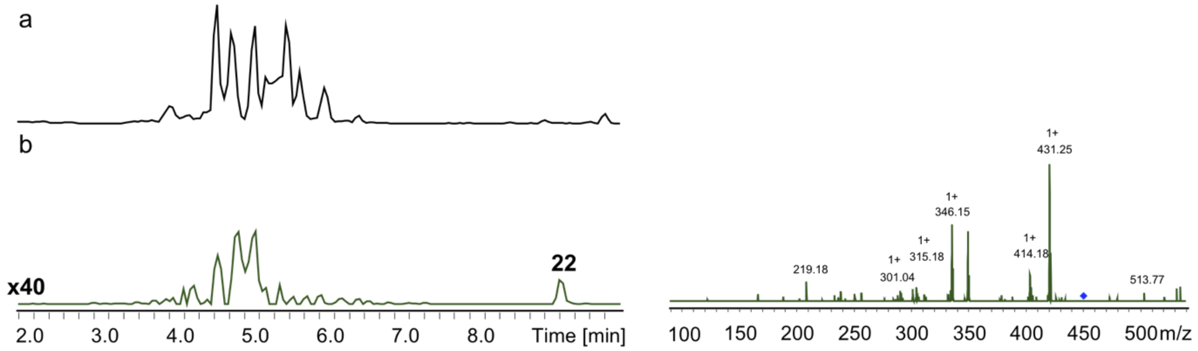

**Figure S23.** HPLC/MS data refers to Figure 6 (NRPS-30) of compound **22** produced in *E. coli* DH10B::*mtaA*. (a) Base Peak Chromatogram (BPC) of an exemplary culture extract. (b) EIC/MS$^2$ data of **22** (*m/z* $[M+H]^+$ = 459.30).

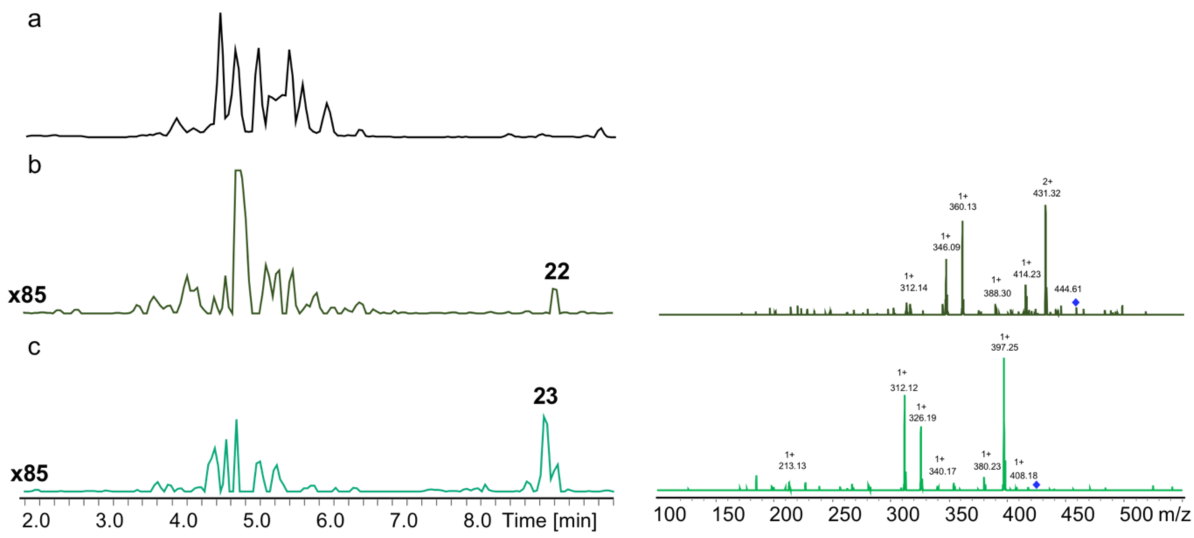

**Figure S24.** HPLC/MS data refers to Figure 6 (NRPS-31) and (NRPS-34) of compounds **22** and **23** produced in *E. coli* DH10B::*mtaA*. (a) Base Peak Chromatogram (BPC) of an exemplary culture extract. (b) EIC/MS$^2$ data of **22** (m/z [M+H]$^+$ = 459.30). (c) EIC/MS$^2$ data of **23** (m/z [M+H]$^+$ = 425.31).

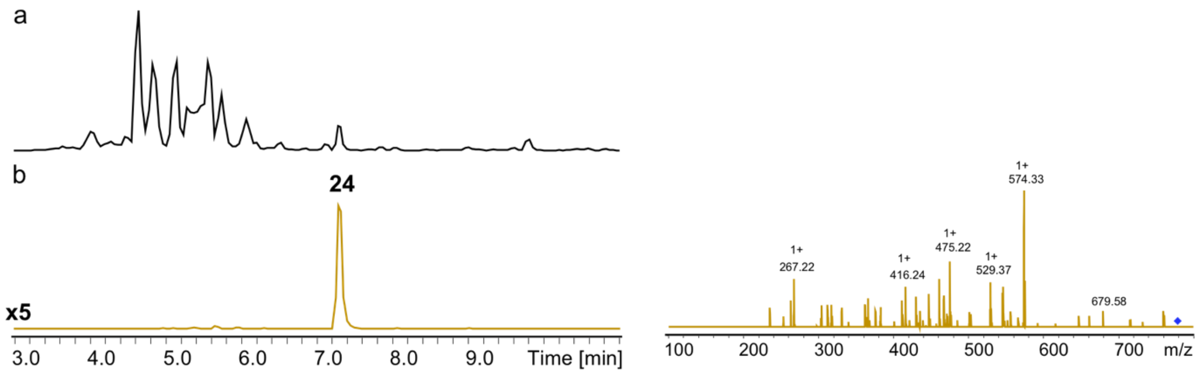

**Figure S25.** HPLC/MS data refers to Figure 6 (NRPS-32) of compound **24** produced in *E. coli* DH10B::*mtaA*. (a) Base Peak Chromatogram (BPC) of an exemplary culture extract. (b) EIC/MS$^2$ data of **24** (*m/z* [M+H]$^+$ = 778.45).

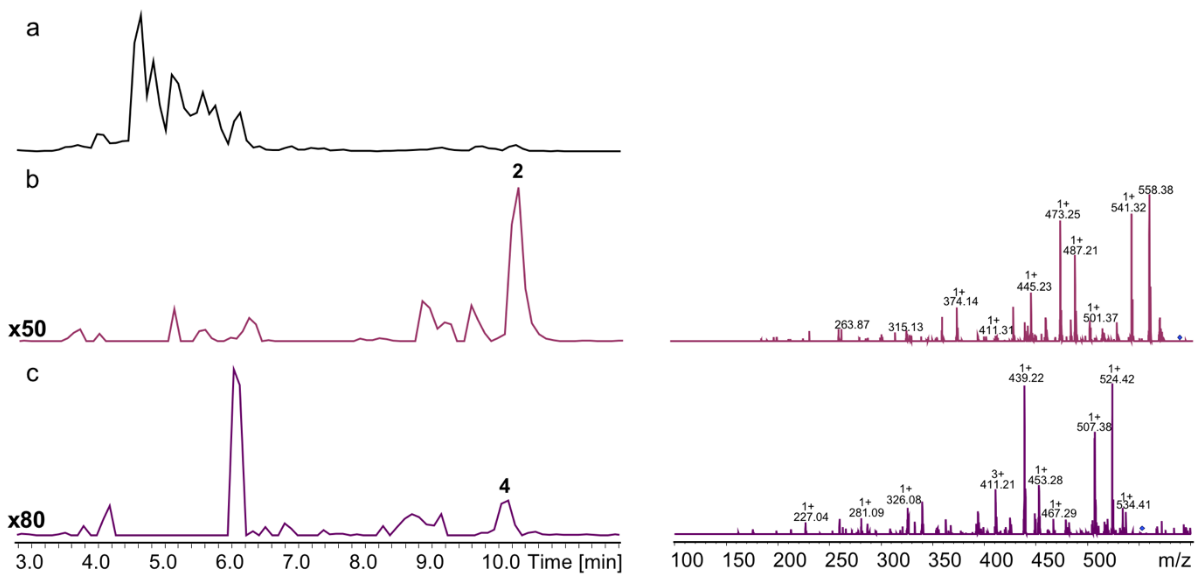

**Figure S26.** HPLC/MS data refers to Figure 6 (NRPS-33) of compounds **2** and **4** produced in *E. coli* DH10B::*mtaA*. (a) Base Peak Chromatogram (BPC) of an exemplary culture extract. (b) EIC/MS$^2$ data of **2** (*m/z* $[M+H]^+$ = 586.40). (c) EIC/MS$^2$ data of **4** (*m/z* $[M+H]^+$ = 552.41).

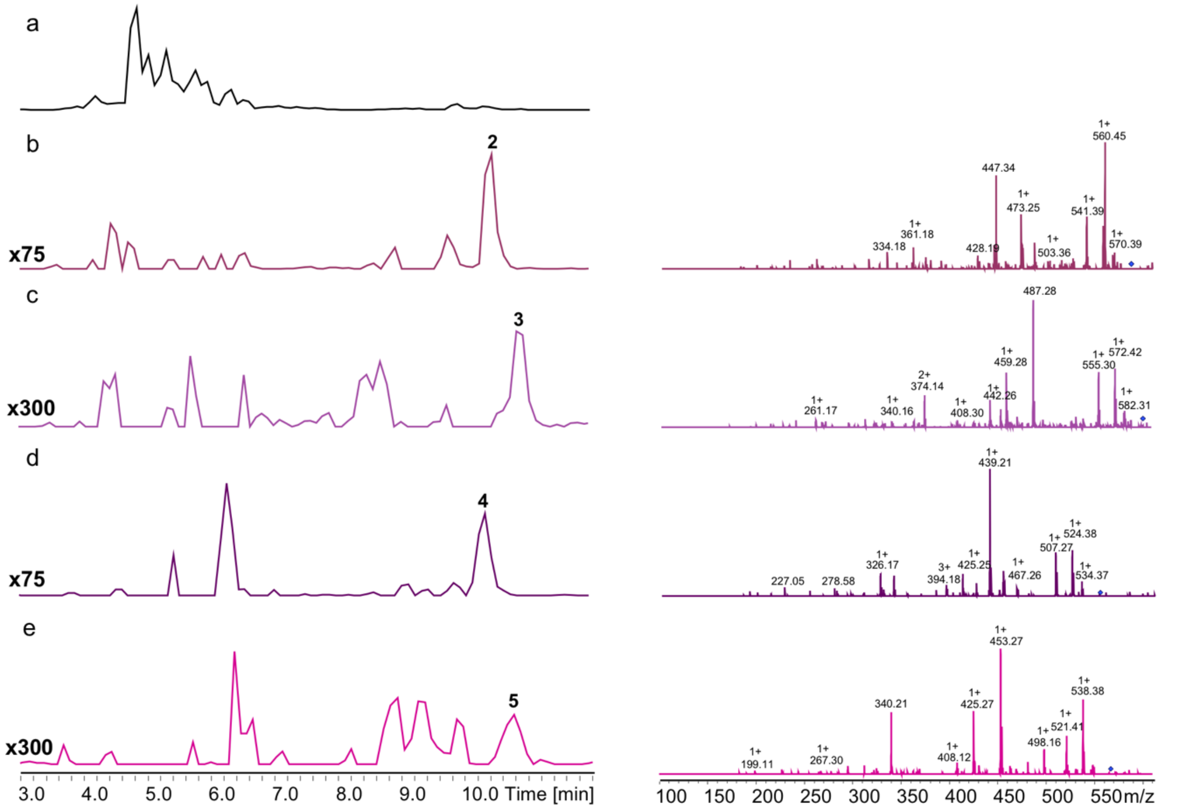

**Figure S27.** HPLC/MS data refers to Figure 6 (NRPS-34) of compounds **2**, **3**, **4** and **5** produced in *E. coli* DH10B::*mtaA*. (a) Base Peak Chromatogram (BPC) of an exemplary culture extract. (b) EIC/MS$^2$ data of **2** (m/z $[M+H]^+$ = 586.40). (c) EIC/MS$^2$ data of **3** (m/z $[M+H]^+$ = 600.41). (d) EIC/MS$^2$ data of **3** (m/z $[M+H]^+$ = 552.41). (e) EIC/MS$^2$ data of **3** (m/z $[M+H]^+$ = 566.43).

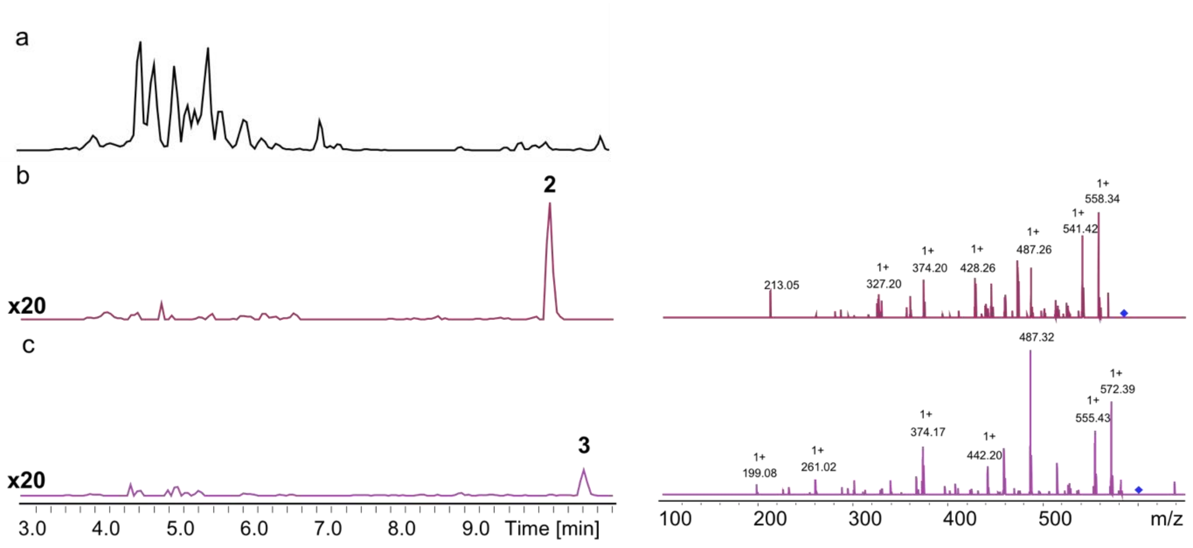

**Figure S28.** HPLC/MS data refers to Figure 6 (NRPS-36) of compounds **2** and **3** produced in *E. coli* DH10B::*mtaA*. (a) Base Peak Chromatogram (BPC) of an exemplary culture extract. (b) EIC/MS$^2$ data of **2** (m/z $[M+H]^+$ = 586.40). (c) EIC/MS$^2$ data of **3** (m/z $[M+H]^+$ = 600.41).

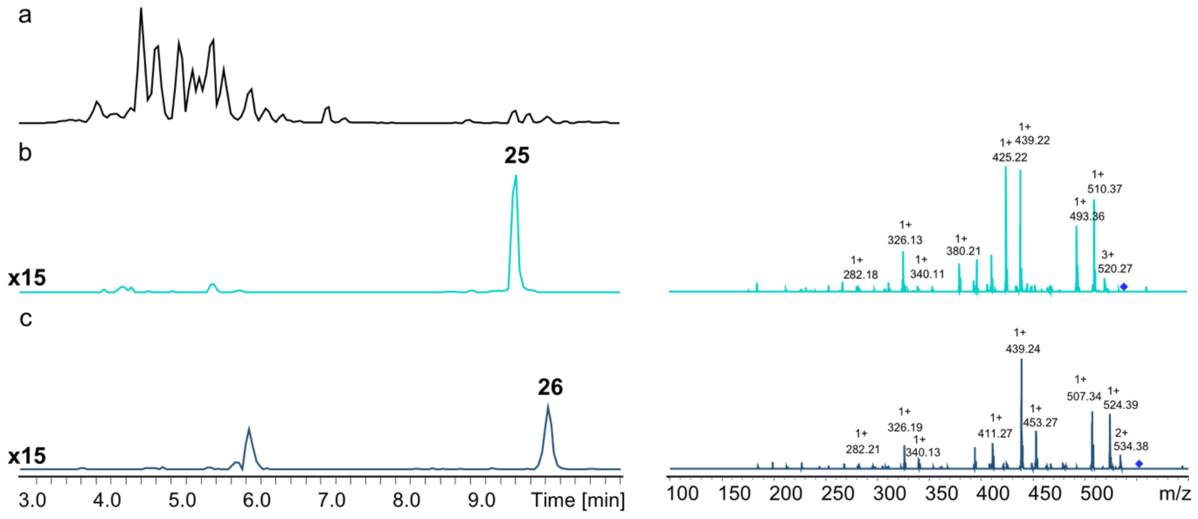

**Figure S29.** HPLC/MS data refers to Figure 6 (NRPS-37) of compounds **25** and **26** produced in *E. coli* DH10B::*mtaA*. (a) Base Peak Chromatogram (BPC) of an exemplary culture extract. (b) EIC/MS$^2$ data of **25** (m/z $[M+H]^+$ = 588.40). (c) EIC/MS$^2$ data of **26** (m/z $[M+H]^+$ = 552.41).

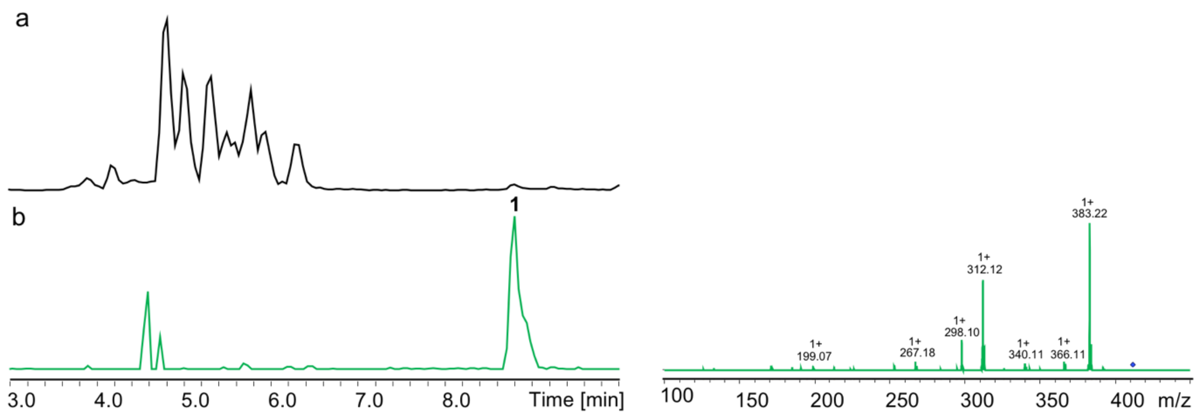

**Figure S30.** HPLC/MS data refers to Figure 6 (NRPS-38) of compound **1** produced in *E. coli* DH10B::*mtaA*. (a) Base Peak Chromatogram (BPC) of an exemplary culture extract. (b) EIC/MS$^2$ data of **1** (m/z $[M+H]^+$ = 411.29).

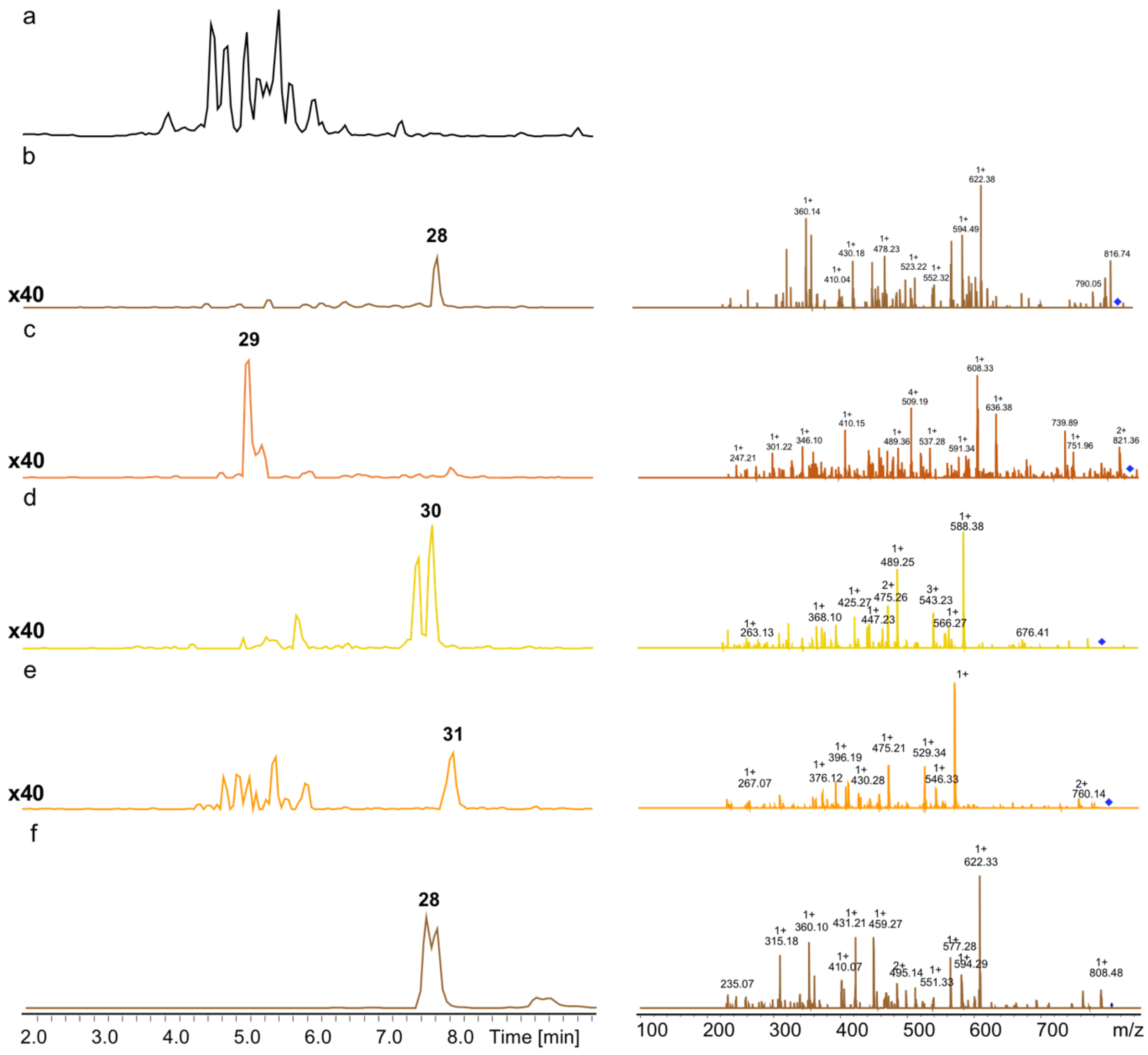

**Figure S31.** HPLC/MS data refers to Figure 6 (NRPS-39) of compounds **28**, **29**, **30** and **31** produced in *E. coli* DH10B::*mtaA*. (a) Base Peak Chromatogram (BPC) of an exemplary culture extract. (b) EIC/MS$^2$ data of **28** (m/z $[M+H]^+$ = 826.45). (c) EIC/MS$^2$ data of **29** (m/z $[M+H]^+$ = 840.47). (d) EIC/MS$^2$ data of **30** (m/z $[M+H]^+$ = 792.47). (e) EIC/MS$^2$ data of **31** (m/z $[M+H]^+$ = 806.48). (f) EIC/MS$^2$ data of synthetic **28** (m/z $[M+H]^+$ = 826.45).

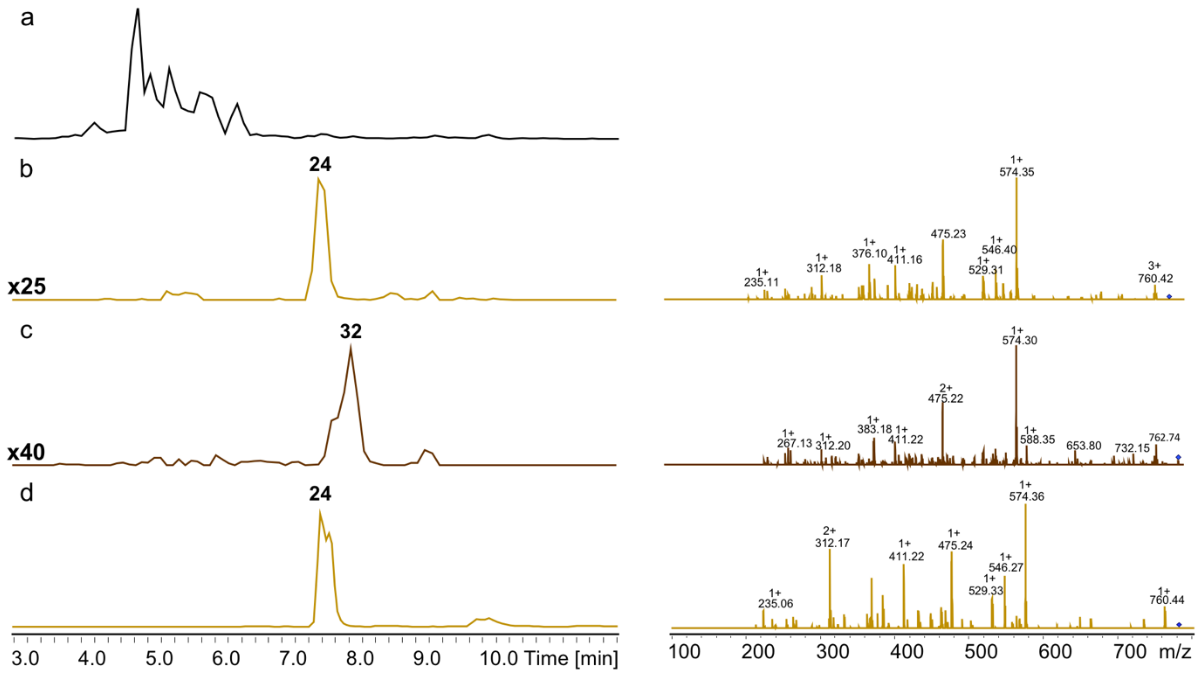

**Figure S32.** HPLC/MS data refers to Figure 6 (NRPS-40) of compounds **24** and **32** produced in *E. coli* DH10B::*mtaA*. (a) Base Peak Chromatogram (BPC) of an exemplary culture extract. (b) EIC/MS$^2$ data of **24** (m/z $[M+H]^+$ = 778.45). (c) EIC/MS$^2$ data of **32** (m/z $[M+H]^+$ = 792.47). (d) EIC/MS$^2$ data of synthetic **24** (m/z $[M+H]^+$ = 778.45).

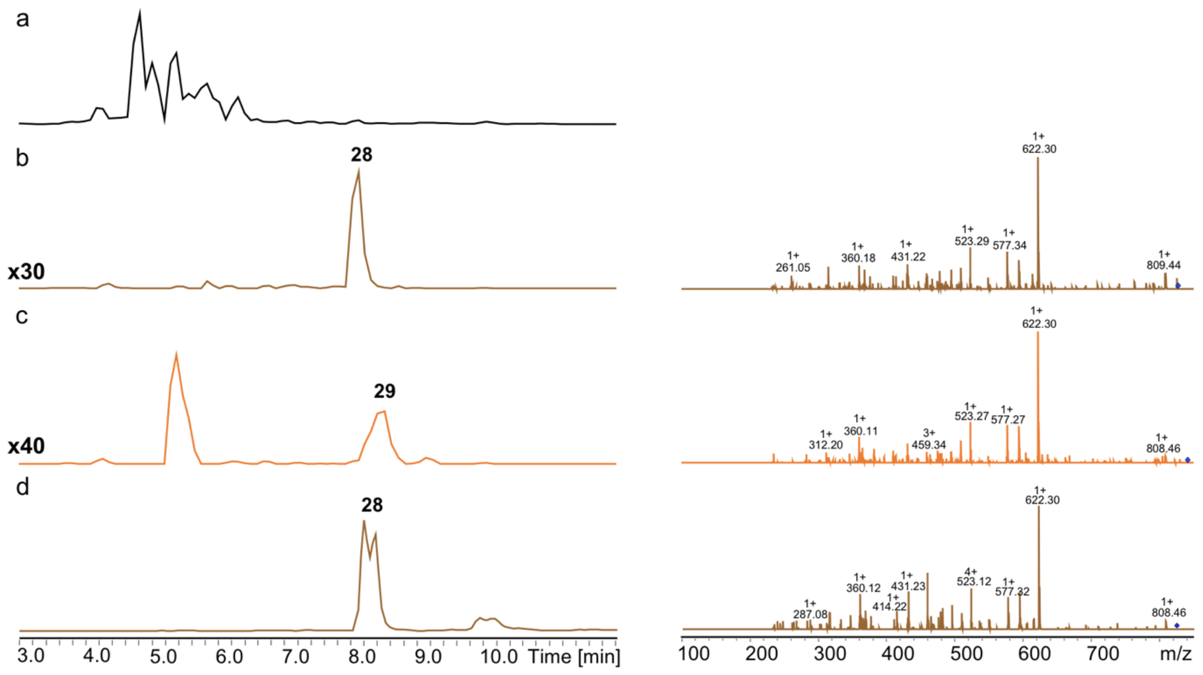

**Figure S33.** HPLC/MS data refers to Figure 6 (NRPS-41) of compounds **28** and **29** produced in *E. coli* DH10B::*mtaA*. (a) Base Peak Chromatogram (BPC) of an exemplary culture extract. (b) EIC/MS$^2$ data of **28** (m/z $[M+H]^+$ = 826.45). (c) EIC/MS$^2$ data of **29** (m/z $[M+H]^+$ = 840.47). (d) EIC/MS$^2$ data of synthetic **28** (m/z $[M+H]^+$ = 826.45).

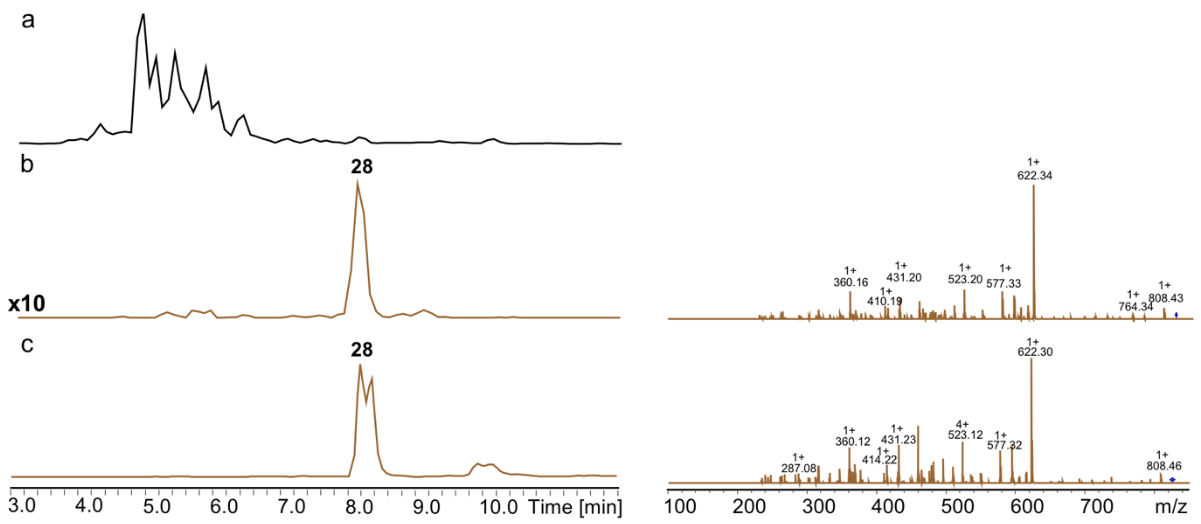

**Figure S34.** HPLC/MS data refers to Figure 6 (NRPS-42) of compound **28** produced in *E. coli* DH10B::*mtaA*. (a) Base Peak Chromatogram (BPC) of an exemplary culture extract. (b) EIC/MS$^2$ data of **28** (m/z $[M+H]^+$ = 826.45). (c) E EIC/MS$^2$ data of synthetic **28** (m/z $[M+H]^+$ = 826.45).

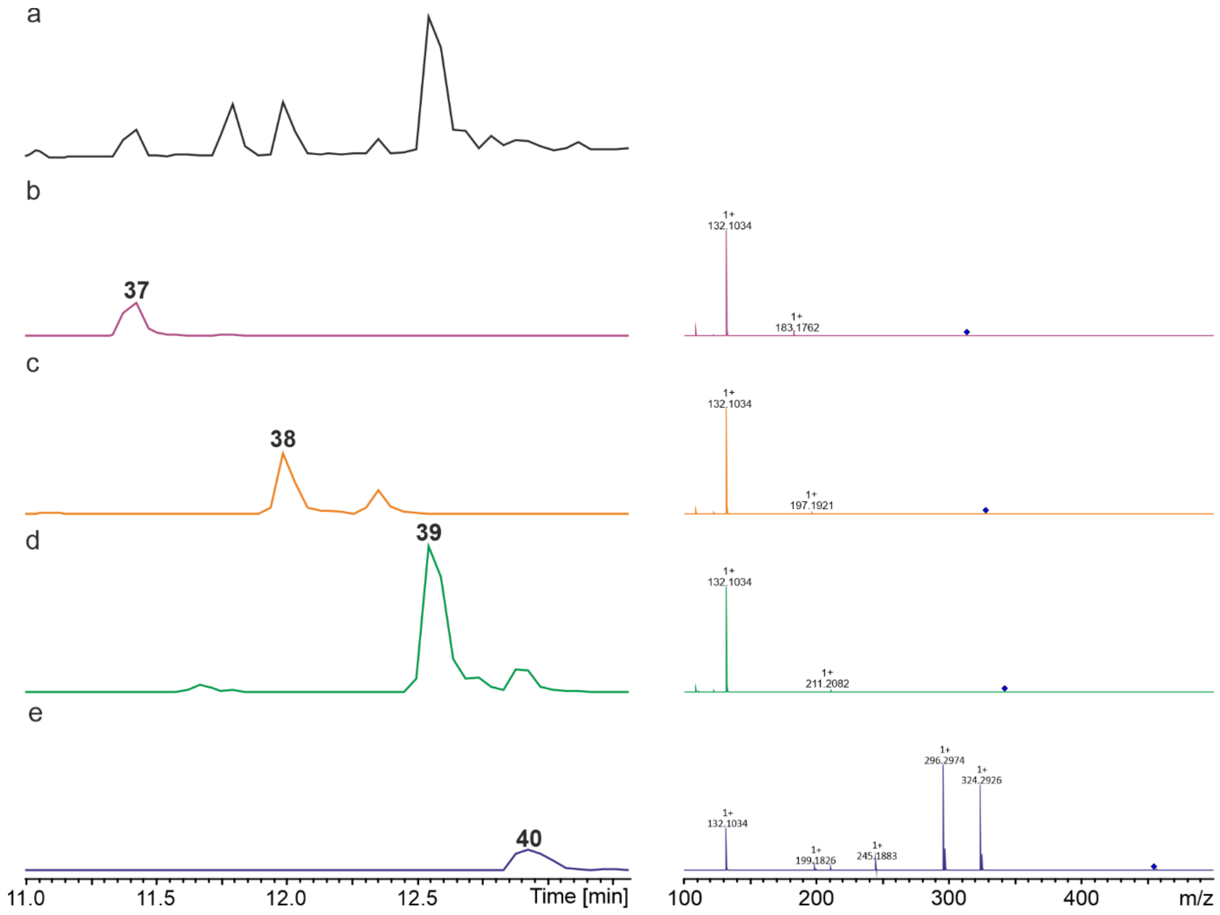

**Figure S35.** HPLC/MS data refers to Supplementary Figure 16 (NRPS-43) of compounds **37**, **38**, **39** and **40** produced in *E. coli* DH10B::*mtaA*. (a) Base Peak Chromatogram (BPC) of an exemplary culture extract. (b) EIC/MS$^2$ data of **37** (m/z $[M+H]^+$ = 314.27). (c) E EIC/MS$^2$ data of **38** (m/z $[M+H]^+$ = 328.29). (d) E EIC/MS$^2$ data of **39** (m/z $[M+H]^+$ = 342.20). (e) E EIC/MS$^2$ data of **40** (m/z $[M+H]^+$ = 455.38).

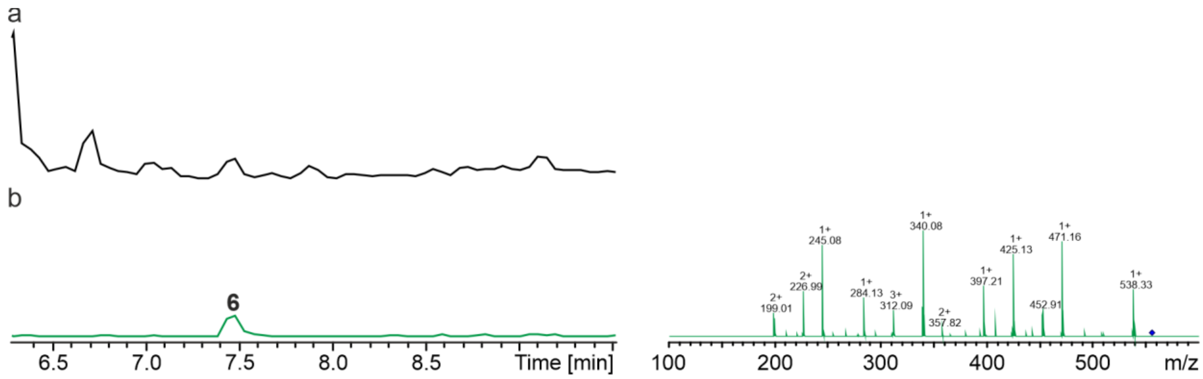

**Figure S36.** HPLC/MS data refers to Supplementary Figure 16 (NRPS-44) of compound **6** produced in *E. coli* DH10B::*mtaA*. (a) Base Peak Chromatogram (BPC) of an exemplary culture extract. (b) EIC/MS$^2$ data of **6** (m/z $[M+H]^+$ = 556.35).

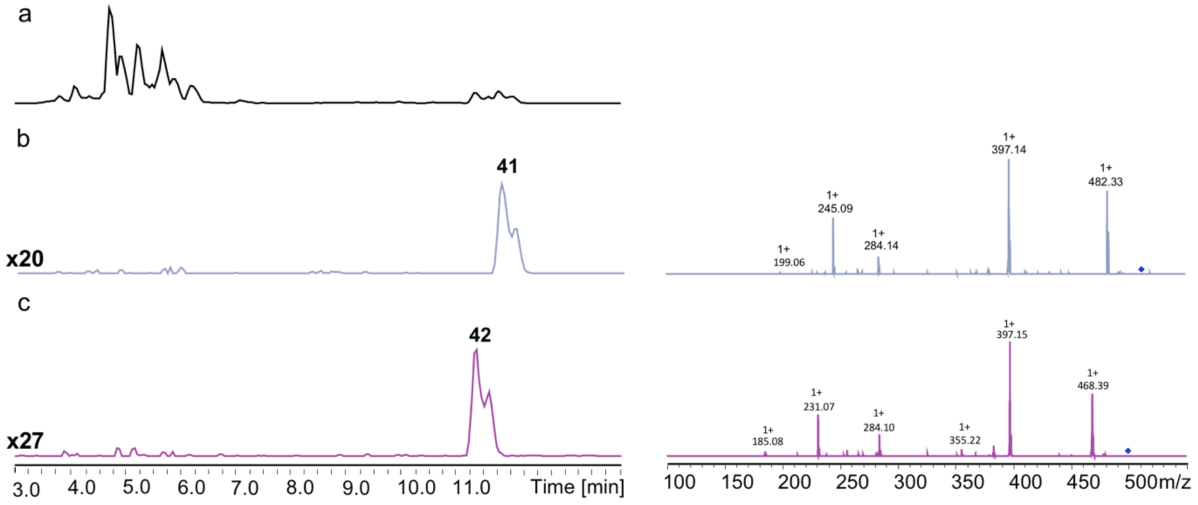

**Figure S37.** HPLC/MS data refers to Supplementary Figure 17 (NRPS-47) of compounds **41** and **42** produced in *E. coli* DH10B::*mtaA*. (a) Base Peak Chromatogram (BPC) of an exemplary culture extract. (b) EIC/MS$^2$ data of **41** (m/z $[M+H]^+$ = 510.39). (c) E EIC/MS$^2$ data of **42** (m/z $[M+H]^+$ = 496.37).

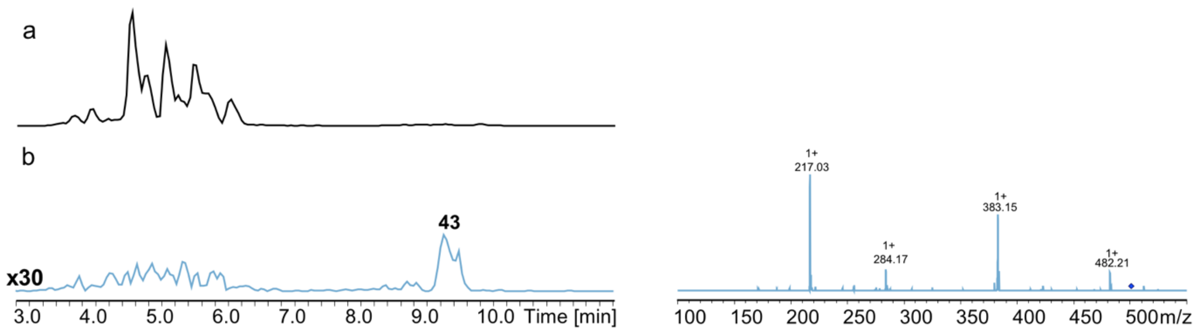

**Figure S38.** HPLC/MS data refers to Supplementary Figure 17 (NRPS-48) of compound **43** produced in *E. coli* DH10B::*mtaA*. (a) Base Peak Chromatogram (BPC) of an exemplary culture extract. (b) EIC/MS$^2$ data of **43** (m/z $[M+H]^+$ = 500.37).
